# A screening strategy based on two zebrafish eleuthero-embryo OECD test guidelines for the hazard assessment of chemicals: case of some bisphenol substitutes

**DOI:** 10.1101/2023.06.16.545329

**Authors:** Armelle Christophe, Benjamin Piccini, Nathalie Hinfray, Edith Chadili, Emmanuelle Maillot-Marechal, Xavier Cousin, Mélanie Blanc, Thierry Charlier, Pascal Pandard, Selim Aït-Aïssa, François Brion

**Author notes:** Corresponding author: François Brion.

## Abstract

The use of efficient screening strategies for the hazard assessment of chemicals is a current challenge to support regulatory requirements. Herein, we combined two eleuthero-embryo assays, a refined Fish Embryo Toxicity assay (OECD TG 236) and the EASZY assay (OECD TG 250), both using transgenic (*tg*) (*cyp19a1b*:GFP). The simultaneous performance of both assays provides complementary information about the acute toxicity, developmental effects, and estrogenic activity. A refined EASZY assay is however necessary to obtain accurate EC50. In this work we compared bisphenol A (BPA) and ten of its substitutes. In the refined FET, most of the selected bisphenols were more toxic than BPA, induced developmental effects on zebrafish embryos, some being identified as teratogenic compounds (BPF, BPS-MAE, BPC Cl, 4,4’ODP), and ten of them induced GFP intensity. Endocrine activity of the BPs was further investigated in the EASZY assay at concentrations that do not affect the survival and the hatching rates or induce developmental toxicity based on the target concentrations used as previously defined in the refined FET. All bisphenols elicited an estrogenic activity with the notable exception of TCBPA. Most BPs were more estrogenic than BPA, acted as agonist ligands of zfERβ2 as shown in zebrafish-specific in vitro reporter gene assay and functional zfERs were required to induce brain aromatase. Interestingly, BPS-MAE and BPS-MPE behave as pro-estrogens as they were unable to transactivate zebrafish ERβ2 *in vitro* but induced brain aromatase *in vivo*. Overall, the implementation of the zebrafish eleuthero embryo-based screening strategy efficiently provided relevant data contributing to their environmental hazard. It also provides further evidence that bisphenols modulate *cyp19a1b* expression during early brain development whom potential short and long-term adverse effects need to be addressed.

**SYNOPSIS:** a zebrafish eleuthero embryo-screening strategy based on OECD TGs was implemented for an efficient hazard assessment of bisphenols revealing that most of them are more toxic and/or estrogenic than BPA

## INTRODUCTION

Bisphenol A (BPA) is an endocrine disrupting chemical (EDC) which interaction with nuclear receptors is well established in various models and thereby causes adverse health effects in human and wildlife. As a result, the use of BPA in food contact materials has been restricted or banned in several countries and the European Union recognized BPA as an EDC for humans and the environment (Rochester, 2013; Oehlmann et al., 2009). Other bisphenols (BPs), structurally or functionally related to BPA, intended to replace it for various industrial applications are increasingly used and detected in humans and in the environment (Chen et al. 2016; Sun et al. 2017; Yu et al. 2015). Some of these substitutes, such as BPF, BPS or BPB, have already been investigated revealing their endocrine activity (Molina et al., 2013, Le Fol et al., 2017) and their reproductive, metabolic, and developmental effects in various models, thereby raising concerns about their health effects in human and wildlife and stimulating the regulatory needs to evaluate their endocrine properties (Serra et al., 2019).

However, for most of BPs, information on their potential endocrine activity is lacking or only partially available, preventing assessment of risks they could potentially posed to human and wildlife. In that perspective, the implementation of efficient screening and testing strategies for the health and environmental hazard assessment of the diversity of substances that are used as BPA substitutes represents a major current challenge.

During the past decade, the zebrafish embryo has become an essential alternative vertebrate model to animal testing in (eco)toxicology studies. At the regulatory level, two zebrafish eleuthero-embryo assays have been adopted as test guidelines (TG) at the OECD level, one to assess the acute toxicity of chemicals, i.e. the zebrafish embryo toxicity test (FET) (TG N° 236) (OECD 2013) and one to assess the estrogenic activity of test chemicals at non-toxic concentrations, i.e., the EASZY assay which was recently adopted (TG N°250) (OECD, 2021).

EASZY is the very first screening assay designed to inform on the estrogenic activity of chemicals acting through estrogen receptors (ERs) of transgenic *tg(cyp19a1b:GFP)* zebrafish. It is also the only OECD test guideline allowing to assess the endocrine activity of test chemicals within the developing brain in a vertebrate model.

The transgenic line *tg(cyp19a1b-GFP)* as proven to reliably informs the acute and developmental effects of test chemicals as compared to wildtype zebrafish embryos usually used in FET assay (Christophe et al., in revision). Herein, we aimed at assessing the toxicity, developmental effect, and estrogenic activity of BPA and ten substitutes by using a zebrafish embryo screening strategy based on *tg(cyp19a1b-GFP)* line and combining both a refined FET assay and the EASZY assay. This testing strategy was complemented using a zebrafish-specific *in vitro* reporter gene assay (Cosnefroy et al., 2012) to gain additional mechanistic information on the ability of BPA substitutes to bind to and activate zfERβ2, the first of the three ER subtypes to emerge within the brain during embryonic development (Mouriec et al., 2009). The data collected are discussed with respect to existing available data and potential regulatory outcomes.

## MATERIALS AND METHODS

### Zebrafish husbandry

Sexually mature transgenic *tg(cyp19a1b:GFP)* zebrafish (*Danio rerio)* were used as breeding stocks. The fish were raised in a recirculating water system (Techniplast, France) at 27°C under a controlled photoperiod (14 h light/10 h dark cycle). They were fed twice a day (dry food and freshly hatched *Artemia salina* nauplii). For breeding, a spawning tray was placed in each aquarium with a sex ratio of 2:1 (male:female), and spawning was stimulated by light. Eggs were collected and cleaned the following morning.

### Chemicals and reagents

All the bisphenols were purchased from Sigma-Aldrich (Saint-Quentin Fallavier, France) except TCBPA (Artmolecule, France), BPS-MPE (Santa Cruz Biotechnology), 4,4’-ODP (TCI Europe n.v.) and BPS-MAE (Ark Pharm. Inc.). All relevant information about selected chemicals is summarized in **Table 1**. Chemicals were dissolved in dimethyl sulfoxide (DMSO, purity 99.5%, Sigma-Aldrich). Reconstituted water (ISO 6341) was prepared every week for controls and preparation of testing chemicals.

**Table 1.**
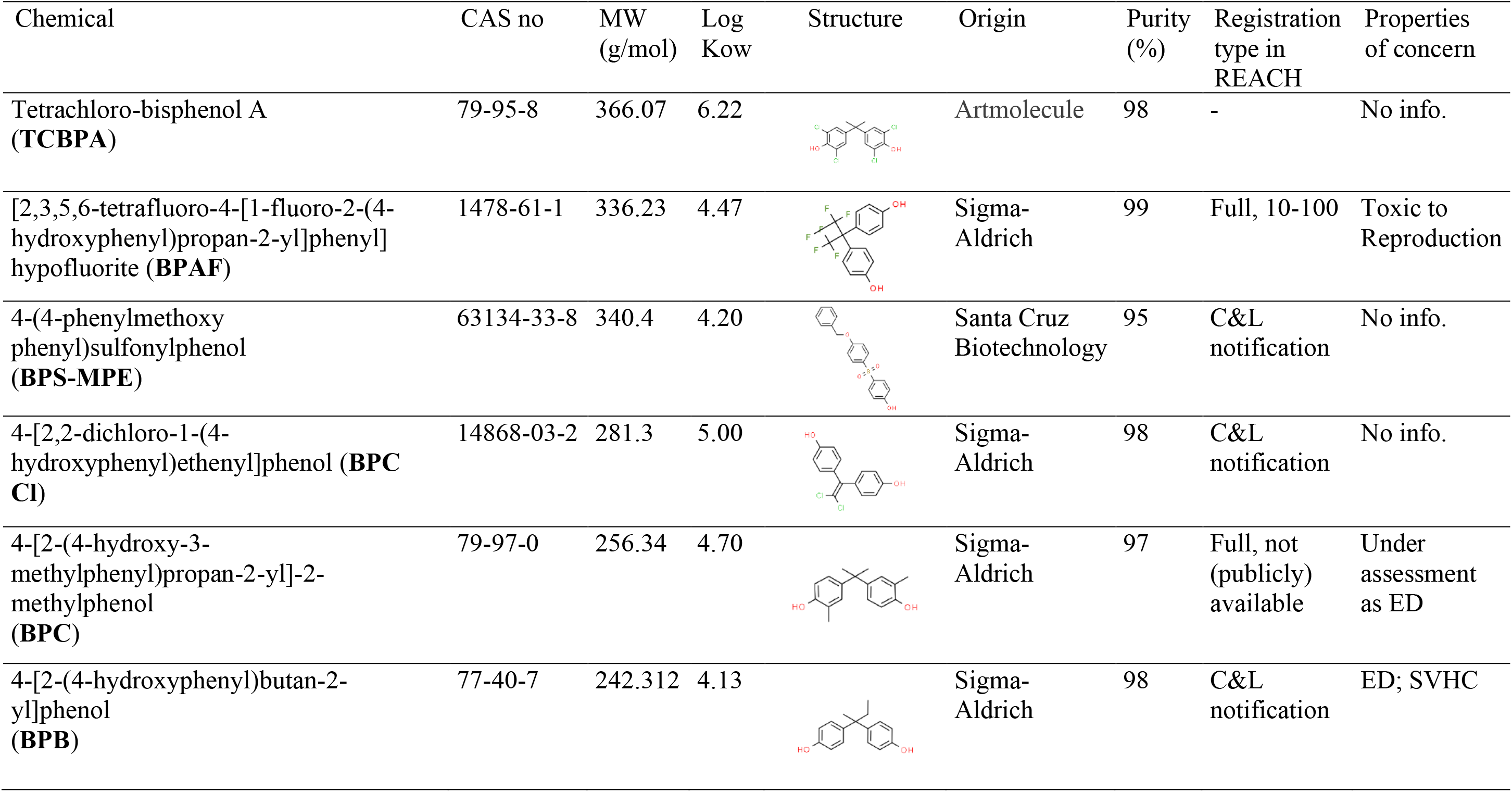
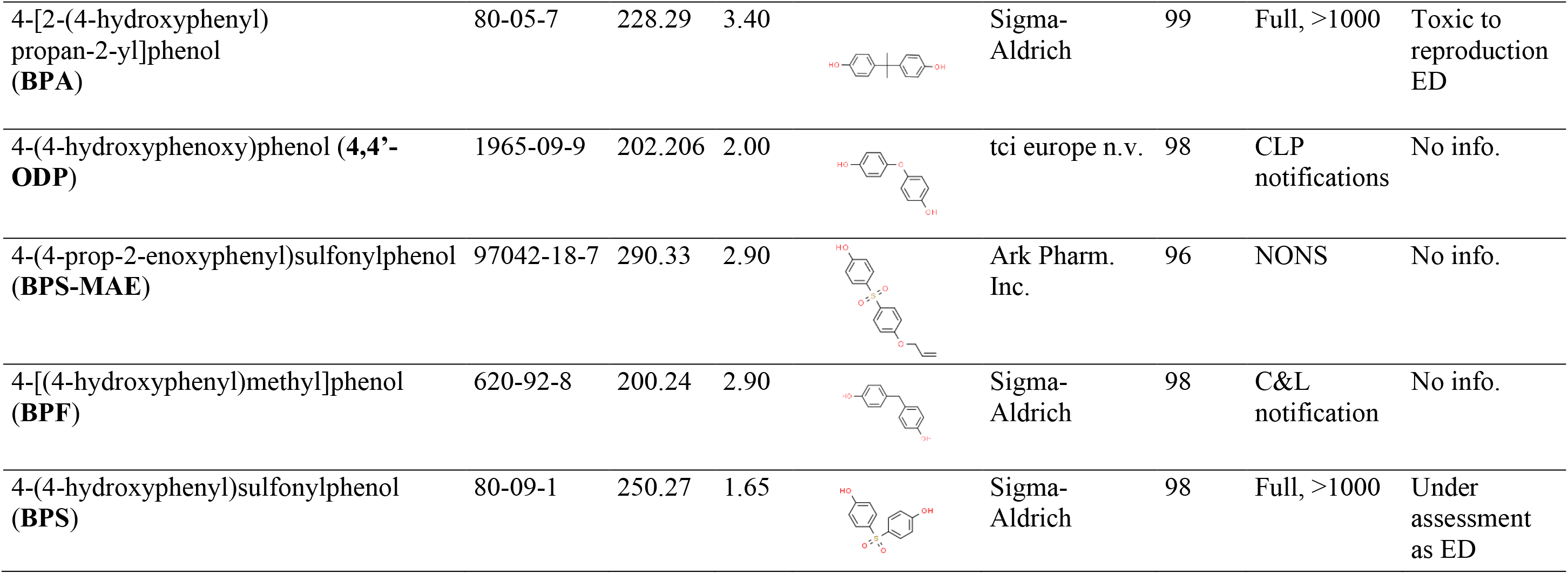
List of bisphenols tested.

| Chemical | CAS no | MW<br>(g/mol) | Log<br>Kow | Structure | Origin | Purity<br>(%) | Registration<br>type in<br>REACH | Properties<br>of concern |
| --- | --- | --- | --- | --- | --- | --- | --- | --- |
| Tetrachloro-bisphenol A<br>(**TCBPA**) | 79-95-8 | 366.07 | 6.22 |  | Artmolecule | 98 | - | No info. |
| [2,3,5,6-tetrafluoro-4-[1-fluoro-2-(4-hydroxyphenyl)propan-2-yl]phenyl]hypofluorite (**BPAF**) | 1478-61-1 | 336.23 | 4.47 |  | Sigma-Aldrich | 99 | Full, 10-100 | Toxic to Reproduction |
| 4-(4-phenylmethoxyphenyl)sulfonylphenol (**BPS-MPE**) | 63134-33-8 | 340.4 | 4.20 |  | Santa Cruz Biotechnology | 95 | C&L notification | No info. |
| 4-[2,2-dichloro-1-(4-hydroxyphenyl)ethenyl]phenol (**BPC Cl**) | 14868-03-2 | 281.3 | 5.00 |  | Sigma-Aldrich | 98 | C&L notification | No info. |
| 4-[2-(4-hydroxy-3-methylphenyl)propan-2-yl]-2-methylphenol (**BPC**) | 79-97-0 | 256.34 | 4.70 |  | Sigma-Aldrich | 97 | Full, not (publicly) available | Under assessment as ED |
| 4-[2-(4-hydroxyphenyl)butan-2-yl]phenol (**BPB**) | 77-40-7 | 242.312 | 4.13 |  | Sigma-Aldrich | 98 | C&L notification | ED; SVHC |
| 4-[2-(4-hydroxyphenyl)propan-2-yl]phenol (**BPA**) | 80-05-7 | 228.29 | 3.40 |  | Sigma-Aldrich | 99 | Full, >1000 | Toxic to reproduction ED |
| 4-(4-hydroxyphenoxy)phenol (**4,4'-ODP**) | 1965-09-9 | 202.206 | 2.00 |  | tci europe n.v. | 98 | CLP notifications | No info. |
| 4-(4-prop-2-enoxyphenyl)sulfonylphenol (**BPS-MAE**) | 97042-18-7 | 290.33 | 2.90 |  | Ark Pharm. Inc. | 96 | NONS | No info. |
| 4-[(4-hydroxyphenyl)methyl]phenol (**BPF**) | 620-92-8 | 200.24 | 2.90 |  | Sigma-Aldrich | 98 | C&L notification | No info. |
| 4-(4-hydroxyphenyl)sulfonylphenol (**BPS**) | 80-09-1 | 250.27 | 1.65 |  | Sigma-Aldrich | 98 | Full, >1000 | Under assessment as ED |

### Refined Fish embryo acute toxicity (FET) tests using *tg(cyp19a1b-GFP)* zebrafish embryos

All the tests were conducted according to the OECD TG 236 (OCDE 2013). The concentration ranges tested, and percentage of solvent used are recapitulated in **Table S1**. During the time course of the refined FET tests, embryos were observed at 8 hours post-fertilization (hpf) (coagulation), 24 hpf (lack of somite formation and non-detachment of the tail) and at 48, 72 and 96 hpf (non-detection of the heartbeat). Based on these criteria, the cumulative mortality rate was calculated at 96 hours, and lethal concentrations inducing 50% mortality (LC50) were derived. The hatching rate was monitored daily. Non-lethal developmental effects were also recorded as previously described (Christophe et al. 2022). At the end of exposure (96 hours), living transgenic zebrafish embryos from test concentrations which do not induce more that 10% of mortality were collected for fluorescence imaging in order to acquire preliminary information about the potential estrogenic activity of the test chemicals. To do so, each embryo was photographed and analyzed as described by Brion et al. (2012).

### EASZY assay

The tests were conducted according to the OECD TG 250 (OCDE 2021). Newly fertilized *cyp19a1b*-GFP eggs were transferred to crystallizers and exposed from 0 to 4 days post-fertlization (dpf) in an incubator (28 ± 1°C) under semi-static conditions with a total renewal of the test solutions each day. Each condition consisted of a triplicate of crystallizers each containing 7 embryos and 15 mL of test medium. Embryos were exposed to a range of non-lethal concentrations of chemicals as determined previously based on the refined FET assays. The tested concentrations range of bisphenols are summarized in **Table S2**. At the end of exposure, transgenic zebrafish embryos were collected for fluorescence imaging to subsequently quantify the GFP intensity as described in the OECD TG N°250 (OECD, 2021).

### In vitro assay

ZELH-zfERs reporter cell line was previously established after a two-step stable transfection of the zebrafish hepatic cell line ZFL with the luciferase gene under the control of ERE (yielding the ZELH cell line that expresses no functional ER) and the zebrafish estrogen receptor subtype β2 (yielding the ZELH-zfERβ2 cell line) (Cosnefroy et al 2012). Cells were cultured, exposed to chemicals for 72 h and measured for luciferase activity exactly as previously described (Cosnefroy et al. 2012, Le Fol et al 2017). For each chemical, serial dilutions were prepared in DMSO and tested in three independent experiments. Final DMSO concentration in the culture medium was always 0.1 % v/v.

### Data analysis

Concentration–mortality curves were modelled using the Regtox 7.5 Microsoft ExcelTM macro which uses the Hill model (http://www.normalesup.org/~vindimian/fr_download.html) to derive lethal concentration inducing 50% of mortality (LC50 values) at 96 hrs with 95% confidence interval (95% CI). For each treatment, the lowest-observed effect concentration (LOEC) was determined for each malformation criterion observed if at least 10% of individuals were affected and if the parameter exhibited a concentration- and/or time-dependent response. Based on these observations, the cumulative percentage of individuals that were malformed was calculated and concentration–response curves for malformation were modeled to derive effective concentration leading to 50% of malformed embryos (EC50). A teratogenic index (TI) was calculated as the ratio LC50/EC50

For fluorescence imaging, GFP fold induction was expressed as mean ± SEM and values were compared to DMSO control using one-way ANOVA followed by Dunnett post-hoc test. For cell exposure, results were expressed as % relative to the maximal effect induced by the positive control 17β-estradiol (E2, 10 nM) and concentration-response curves were modeled using the RegTox Excel macro to derive EC50 values. In the case of incomplete concentration-response curves (*i.e.*, no plateau observed), the maximal effect parameter was fixed to 100% activation and a PC50 (concentration inducing 50% of the maximal positive control response) value was determined.

## RESULTS AND DISCUSSION

### Most of the selected bisphenols tested in the FET assays are more toxic than BPA

Bisphenols induced variable toxicity on *tg(cyp19a1b:GFP)* embryos in the FET tests with LC50(96h) ranging from 0.88 mg/L for TCBPA to 37 mg/L for BPF while BPS was not toxic at the highest tested concentration (**Table 2; Figure S1**). These results allow classifying the BPs according to their acute toxicity: TCBPA > BPAF > BPS-MPE > BPC-Cl > BPC > BPB > BPA > 44’ODP > BPS-MAE > BPF > BPS. For six out of eleven BPs for which previous data were available, our data are in agreement with LC50 reported using wild-type zebrafish embryos (**Table 2, Table S3**), thereby reinforcing the usefulness of *tg(cyp19a1b:GFP)* to provide reliable acute toxicity data in the refined FET test (Christophe et al., under revision). For the five chemicals for which no information was available, namely BPS-MPE, BPC Cl, BPC, 44’ODP and BPS-MAE, this study provides LC50 in the 3.17 to 20.07 mg/L range and reveals that three of them (*i.e.* BPS-MPE, BPC-Cl and BPC) are more toxic than BPA.

**Table 2.**
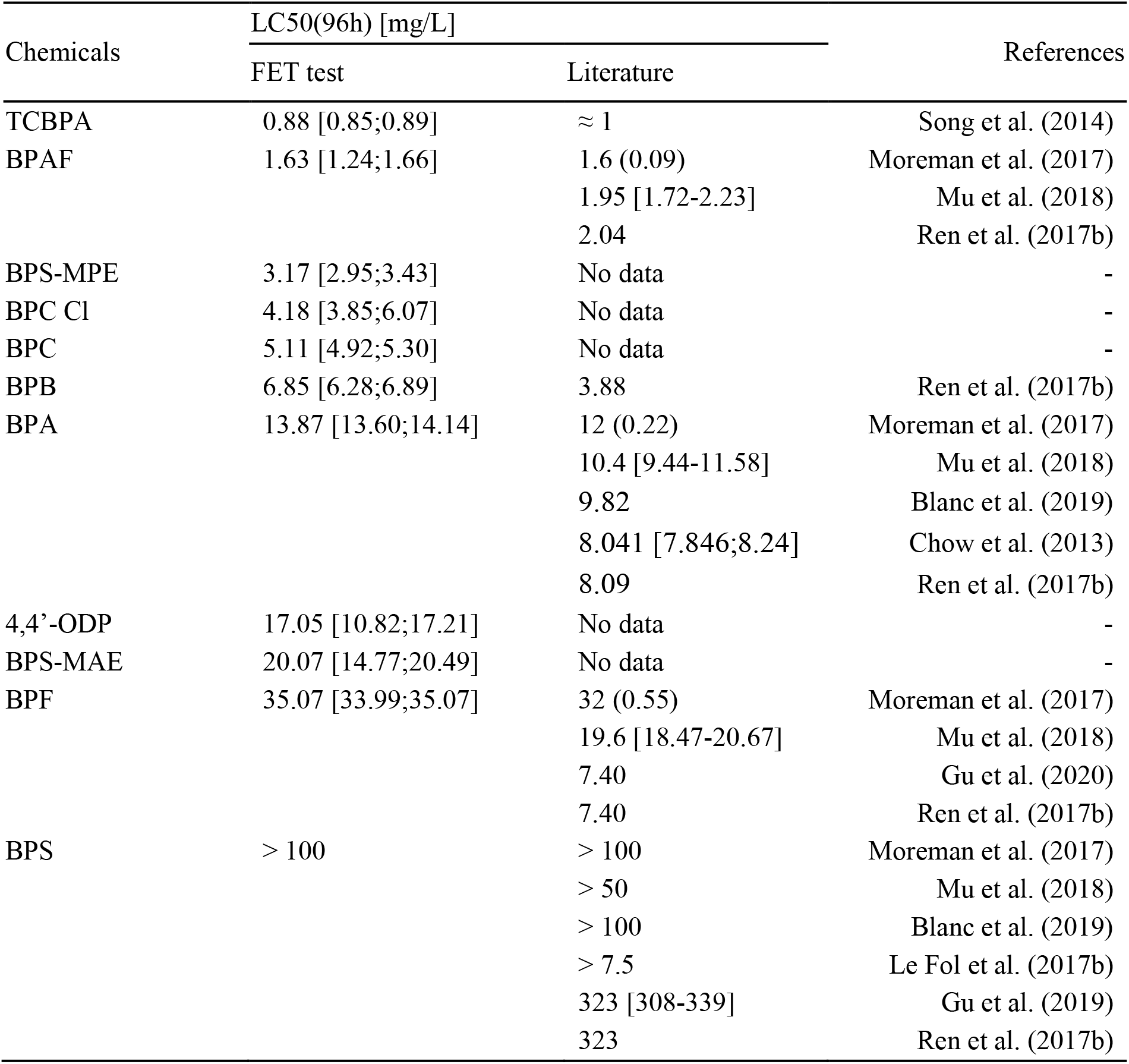
LC50(96h) derived from the refine FET assay for the eleven tested bisphenols. Available LC50(96h) collected from the literature using the FET test are reported when available. Numbers in brackets represent the 95% confidence interval, numbers in parentheses represent SEM.

### Bisphenols induced developmental effects on zebrafish embryos and some are teratogenic

Morphological and physiological changes were also recorded to assess more precisely the potential developmental toxicity of BPs. For most of them a reduced hatching success was observed accompanied by a delayed in hatching time (**Figure S2**). A notable exception was however noted for BPS-MPE and BPS-MAE for which an apparent advanced in hatching time was observed (Figure S2). Furthermore, all the bisphenols studied induced developmental effects, which nature varied depending on the test chemicals and their concentrations of exposure. The most frequently observed phenotypic responses in BPs-exposed zebrafish were deformity of the yolk, edema, heart rate disruption, spinal malformation, and haemorrhage, both of these responses agreeing with those previously reported (**Table S3**). Several BPs elicited similar phenotypic response spectrum suggesting similar mode of action on developmental processes with the notable exception of BPS, which did not show developmental effects except deformity of the yolk observed at the highest concentration tested. While some authors suggested the possibility to assign molecular pathways according to the promoted phenotypic responses (Jarque et al., 2020), a more specific response was reported for six out of the eleven substitutes which were both characterized by a reduction of pigmentation, i.e., BPA, BPF, BPS, BPS-MAE, BPS, MAE, BPC Cl and 4,4’- ODP. Altered pigmentation in zebrafish embryos exposed to BPA and BPF was previously reported in studies highlighting a more pronounced effect for BPF exposures, which agrees with our results (Moreman et al., 2017, Cao et al. 2018; Mu et al. 2020; Oh et al. 2018). In the present study, BPs-induced depigmentation occurred at varying concentrations and intensities depending on the bisphenols tested (**Table 3, Figure S3**). Strikingly, a full depigmentation was observed in 4,4’-ODP-exposed embryos from concentration as low as 0.99 mg/L. This effect likely reflects their ability to disrupt the melanin biosynthetic pathway (Cao et al. 2018; Mu et al. 2020; Oh et al. 2018) with a high efficiency for 4,4’ODP as compared to BPA and BPF.

**Table 3.** LOECs of developmental endpoints observed throughout the FET tests. For each bisphenol, the occurrence of phenotypic or physiological changes was calculated and the EC50 derived (considering all the phenotypic or physiological changes). TI= teratogenic index.

| Endpoints | LOECs (mg/L) |  |  |  |  |  |  |  |  |  |  |
| --- | --- | --- | --- | --- | --- | --- | --- | --- | --- | --- | --- |
|  | TCBPA | BPAF | BPS-MPE | BPC Cl | BPC | BPB | BPA | 4,4'-ODP | BPS-MAE | BPF | BPS |
| Deformity of yolk | 1 | 1 | 5.56 | 0.99 | 2.84 | 10 | 20 | 2.96 | 11.1 | 15 | 400 |
| Growth retardation | - | - | - | - | - | - | 20 | 8.89 | - | - | - |
| No spontaneous movement | 2 | 2 | - | 2.96 | - | 10 | 10 | - | 33.3 | 60 | - |
| Edema | 0.5 | - | 5.56 | 0.99 | 2.84 | 5 | 10 | 0.99 | 33.3 | 15 | - |
| No blood tail circulation | 1 | - | - | 8.89 | 8.53 | - | - | 8.89 | 33.3 | 60 | - |
| Heart rate disruption | 1 | 2 | 5.56 | 8.89 | 8.53 | 10 | 20 | 2.96 | 33.3 | 60 | - |
| No/Less pigmentation | 1 | - | 5.56 | 8.89 | - | - | 10 | 0.99 | 33.3 | 3.75 | - |
| Head malformations | 1 | 2 | - | - | - | 10 | - | 2.96 | - | 7.5 | - |
| Spinal malformations | 0.5 | 1 | 5.56 | 8.89 | 8.53 | - | 10 | 2.96 | 33.3 | 30 | - |
| Hemorrhage | - | 1 | 5.56 | 2.96 | 8.53 | 10 | 10 | 0.99 | - | 7.5 | - |
| EC50 (mg/L) (cumulative sublethal endpoints) | 0.41 | 1.07 | 100 | 1.12 | 2.08 | 3.03 | 6.06 | 0.50 | 6.41 | 1.56 | 263 |
| TI | 2.1 | 1.5 | 0.03 | 3.7 | 2.5 | 2.3 | 2.3 | 34.4 | 3.1 | 22.5 | 0.4 |

When endpoints are considered as relevant, lowest-observed effect concentrations (LOECs) are raised. Endpoints are considered as relevant when they reach at least 10% of impacted embryos in a condition, and when they are concentration and/or time dependent. EC50 are derived from the concentration-response curves established for percentage of affected embryos at 96 hpf for each concentration tested for the 11 bisphenols. With these EC50 and with LC50 obtained in the FET tests, we calculated a relative teratogenic index (TI) for each chemical ( $TI = LC50/EC50$ ).

Numerous studies reported zebrafish embryos as a useful model to identify (pro)teratogenic compounds and to predict levels of potential outcomes in mammals with high accuracy using relative teratogenic index (TI) calculation (Selderslaghs et al., 2009, Weigt et al., 2011, Jarque et al., 2020). In the present study all BPs were characterized by TI >1 except for BPS and BPS- MPE. Considering that TI ≥ 3 and EC50 <1 mM are suitable limits to differentiate teratogens from non-teratogens in zebrafish (Jarque et al., 2020), we confirmed BPF as a teratogenic compound and newly identified BPS-MAE, BPC Cl and 4,4’ODP as teratogenic bisphenols, thereby contributing to the hazard identification of BP substitutes. Relative teratogenicity of these chemicals is BPS-MAE < BPC Cl << BPF < 4,4’ODP.

### Almost all bisphenols can induce GFP in tg(cyp19a1b:GFP)

We took advantage of using *tg(cyp19a1b:GFP)* embryos in the refined FET assay to image GFP in hatched embryos in order to gain preliminary information about the potential estrogenic activity of tested BPs. Significant inductions of GFP were reported for all the bisphenols with the notable exception of TCBPA for which a two-fold GFP increase was noted as at 0.25 mg/L but not at the higher concentration (**Figure 1, Table S4**). In most cases, the induction profiles did not allow to derive EC50 to quantify their estrogenic activity, expect for BPA and BPS for which full concentration-responses curves were obtained (EC50 = 1.49 mg/L and 43 mg/L for BPA and BPS respectively; **Figure S4, Table S4**).

**Figure 1.**
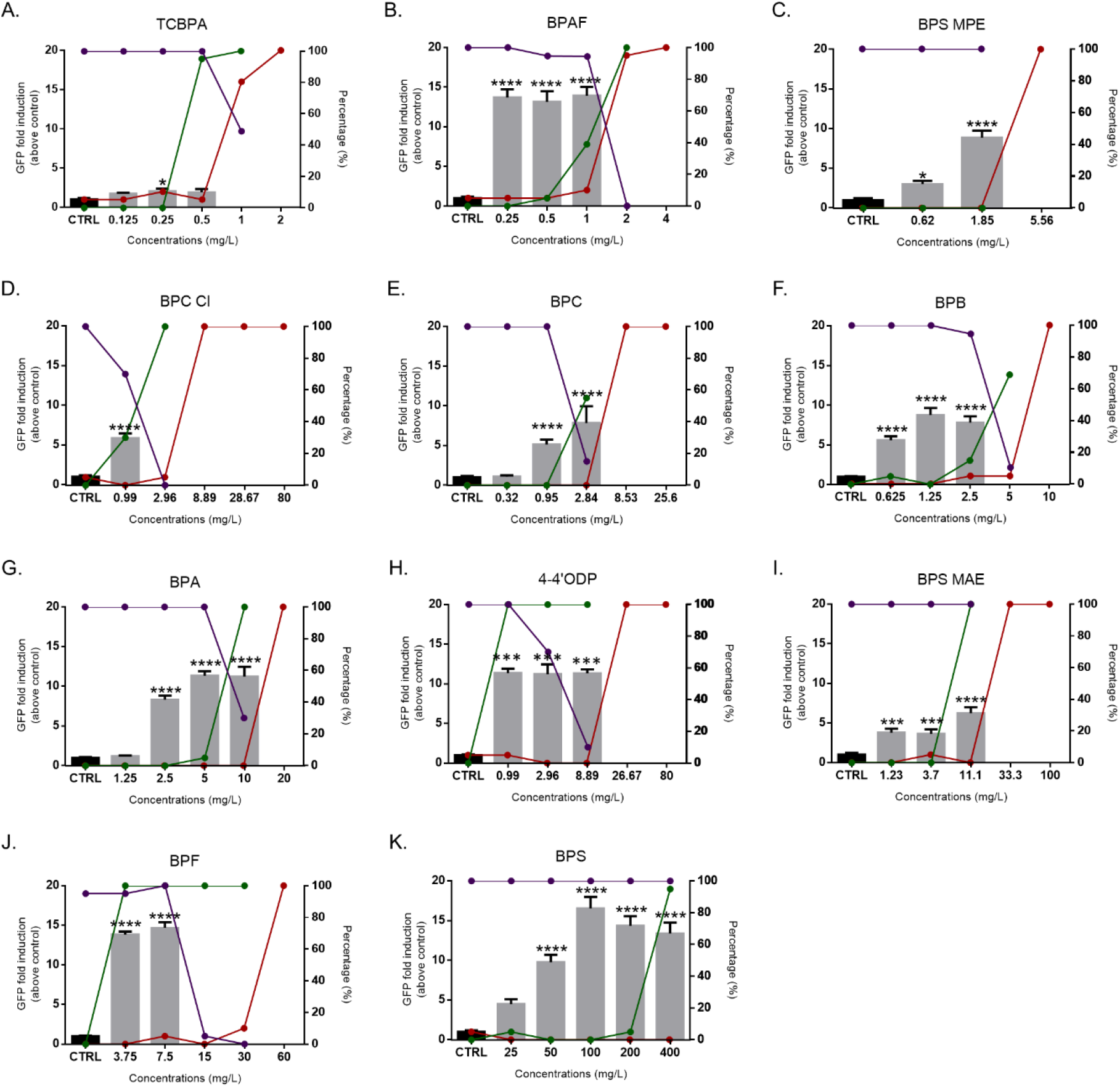
Summary results of the FET tests for each chemical tested. Red lines represent percentage of mortality after 96h of exposure; green lines represent percentage of embryos affected during the exposure; and violet lines represent percentage of hatching at 96h. Histogram represent results of fluorescence imaging at the end of the FET test (96 h), the asterisk symbol denote differences compared with the control group (CTRL = water or solvent control): *p < 0.05, ***p < 0.0005, ****p < 0.0001; data are represented as mean ± SEM. Left axis is the GFP fold induction above control group for fluorescence imaging, and right axis is percentage (%) for mortality, malformed embryos and hatching.

### In the EASZY assay, several bisphenols have higher estrogenic activity as BPA

Based on the preliminary data obtained in the frame of the FET, we refined concentrations to be used in an EASZY assay in order to obtain pore accurate information about estrogenic potential of tested chemicals. Definition of concentrations range to be used was further dictated by three conditions. The maximum concentrations had to fulfil the following requirements: they should not (i) induce mortality, (ii) affect the hatching rate or (iii) significantly induce developmental effects in zebrafish embryos (**Table S3**). Results obtained from the EASZY assays confirmed the absence of estrogenic activity for TCBPA. We also further confirmed the estrogenic activity of all other BPs as shown by the concentration-dependent induction of GFP in the developing brain of zebrafish (**Figure 2**). Different induction profiles were observed (**Figure 3**). Some BPs, i.e., 4,4’-ODP, BPF and BPS, induced full concentration-dependent response curves with maximal response reaching the GFP levels measured in the positive control (EE2 14.8 ng/L) while for other BPs the concentration-response curves were characterized by maximal fold induction levels below GFP levels measured in the positive controls. For BPS-MAE, significant inductions were measured from 0.625 mg/L but unusual concentration-response curve was observed as the response reached a plateau at low induction levels. Such differential induction profiles may reflect, at least partly, the agonist activity of these substances toward zebrafish ERs (see below). Based on these induction profiles, the EC values (**Table 4)** were derived from the modelled concentration-responses curves for GFP (**Figure S5**) to quantify the estrogenic activity of each bisphenol allowing to rank them from the less active: to the most active BPS < BPA ∼ BPS-MPE < BPB < BPC < 4,4’-ODP < BPF < BPS-MAE < BPAF < BPC Cl Estrogenic activities of BPC Cl > BPAF > BPS-MAE > BPF > 4,4’-ODP > BPC and BPB were 54, 13, 2.6, 2.3, 2.2, 1.7 and 1.5-fold higher than by BPA, respectively. BPS-MPE had a similar estrogenic activity but was less potent to stimulate GFP as BPA. Among all the BPs tested, BPS was the only one being less active than BPA. Overall, our study confirms the estrogenic activity of several BPs (i.e., BPAF, BPC, BPA, BPF, BPS) on different zebrafish ER-sensitive embryo models (Moreman et al., 2017, Lefol et al., 2017, Pinto et al., 2019) and provides new quantitative estrogenic activity for five of them (BPC Cl, BPS-MAE, 4,4’-ODP, BPB, BPS-MPE), most of them being characterized by a higher or comparable estrogenic activity than BPA.

**Figure 2.**
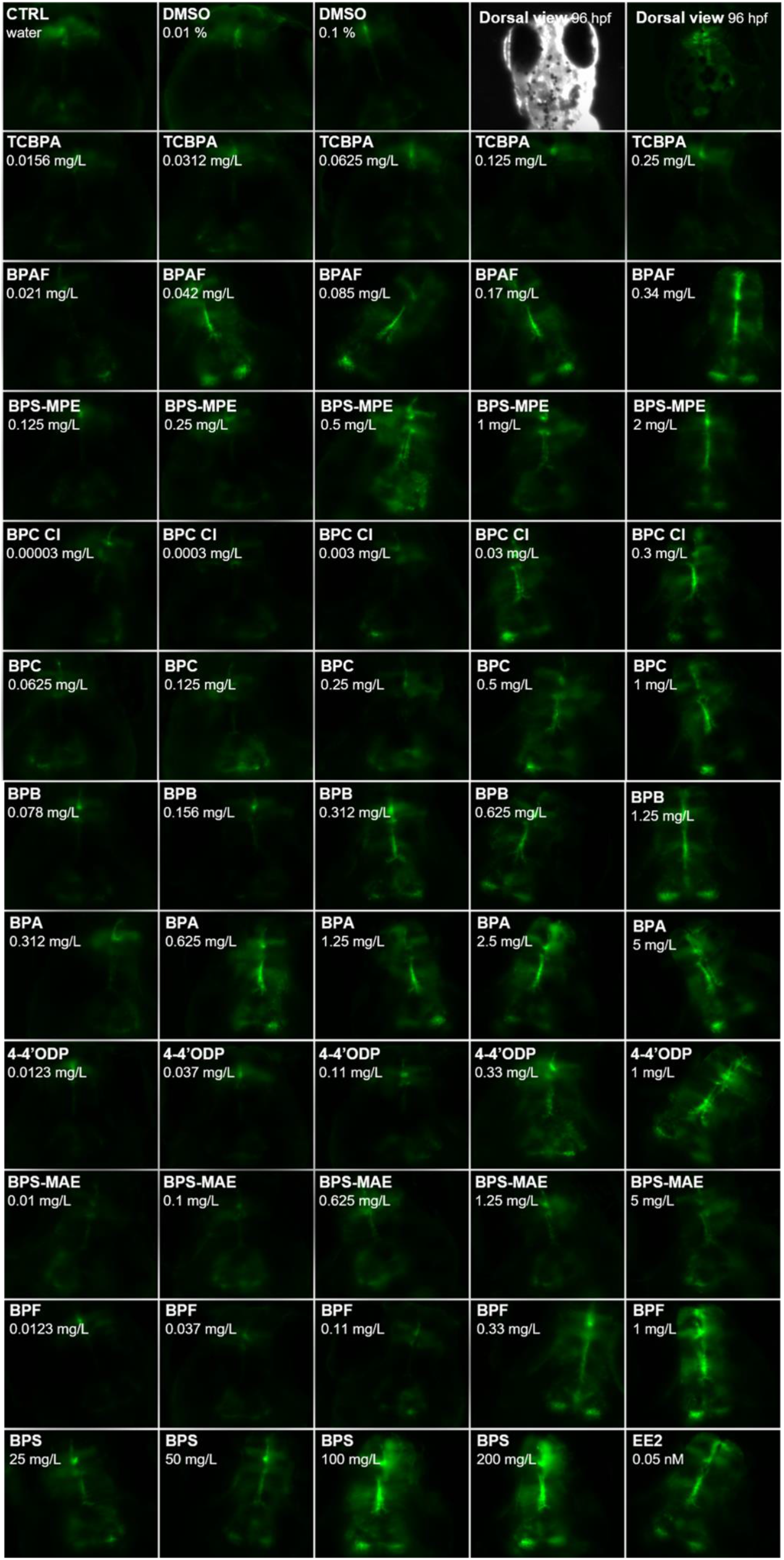
*In vivo* imaging of 96 hpf old live tg(*cyp19a1b*-GFP) zebrafish embryos exposed to chemicals inducing GFP expression in radial glial cells. Dorsal views of zebrafish heads, for each chemical the concentration used is indicated.

**Figure 3.**
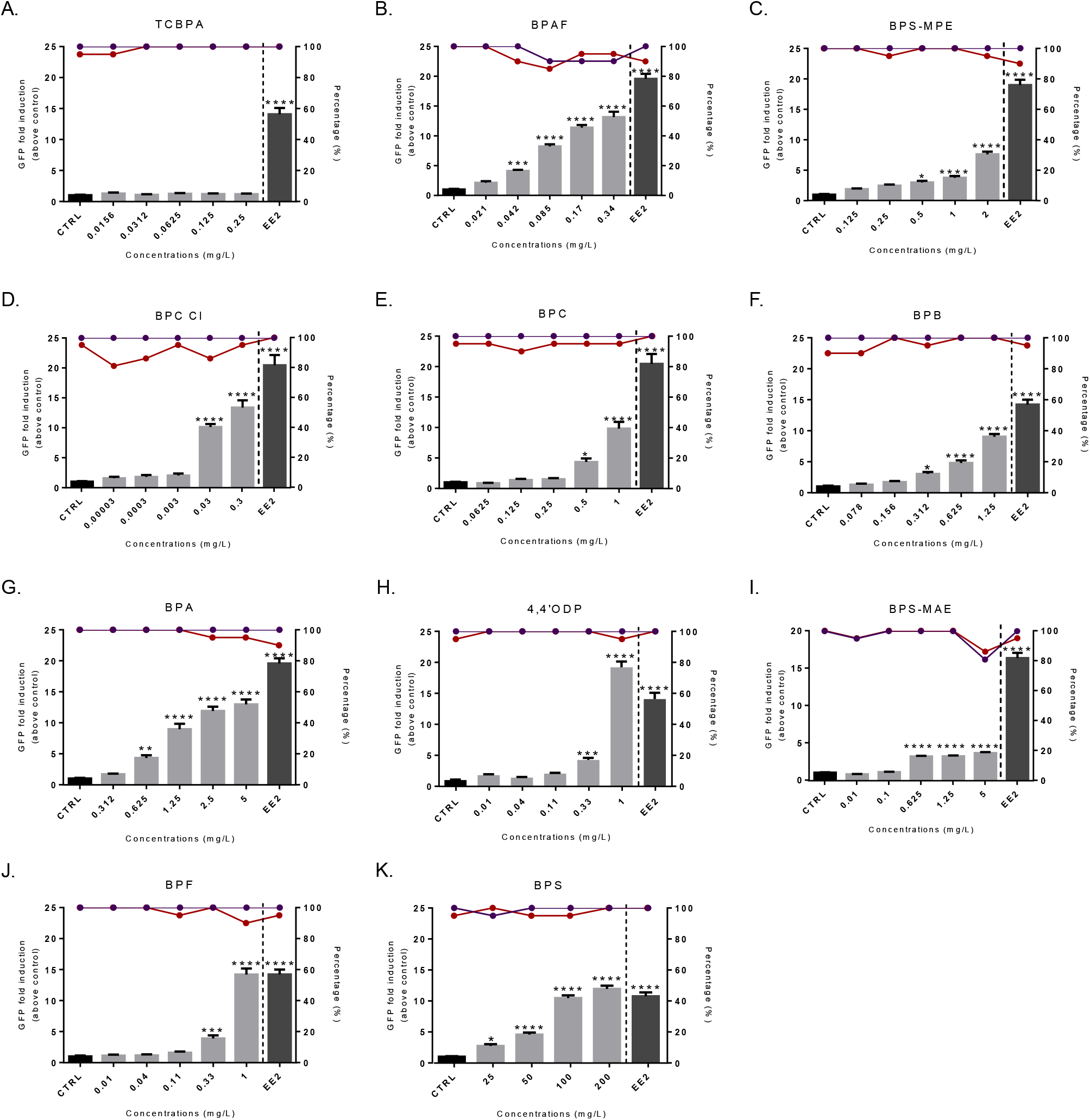
GFP intensity expressed as mean fold induction above control ± S.E.M quantified in *tg(cyp19a1:GFP)* embryos exposed to bisphenols. The red line is the percentage of survival and the violet line is the percentage of hatched embryo after 96h of exposure. the asterisks denote differences compared with the control group (CTRL = solvent control): *p < 0.05, ***p < 0.0005, ****p < 0.0001. Left axis is the GFP fold induction above control group for fluorescence imaging, and right axis is percentage (%) for mortality and hatching.

**Table 4.**
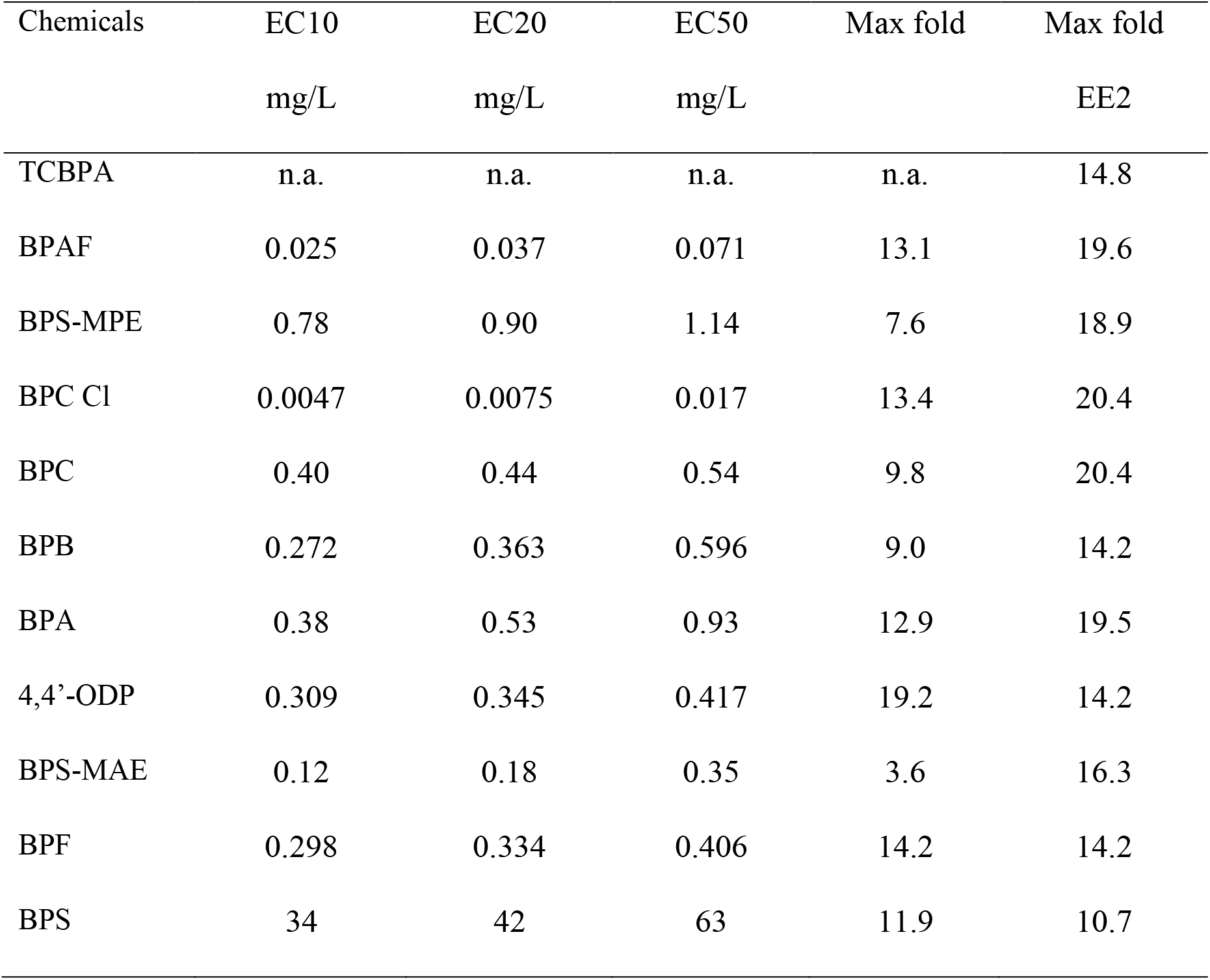
ECx values obtained in the EASZY assays for all the BPs. Data are expressed in mg/L. Max Fold = Mean measured maximum fold induction (Modeled curves used to derived ECx values are shown in figure S5)

| Chemicals | EC10 | EC20 | EC50 | Max fold | Max fold |
| --- | --- | --- | --- | --- | --- |
|  | mg/L | mg/L | mg/L |  | EE2 |
| TCBPA | n.a. | n.a. | n.a. | n.a. | 14.8 |
| BPAF | 0.025 | 0.037 | 0.071 | 13.1 | 19.6 |
| BPS-MPE | 0.78 | 0.90 | 1.14 | 7.6 | 18.9 |
| BPC Cl | 0.0047 | 0.0075 | 0.017 | 13.4 | 20.4 |
| BPC | 0.40 | 0.44 | 0.54 | 9.8 | 20.4 |
| BPB | 0.272 | 0.363 | 0.596 | 9.0 | 14.2 |
| BPA | 0.38 | 0.53 | 0.93 | 12.9 | 19.5 |
| 4,4'-ODP | 0.309 | 0.345 | 0.417 | 19.2 | 14.2 |
| BPS-MAE | 0.12 | 0.18 | 0.35 | 3.6 | 16.3 |
| BPF | 0.298 | 0.334 | 0.406 | 14.2 | 14.2 |
| BPS | 34 | 42 | 63 | 11.9 | 10.7 |

**Table 5.** EC50 values and transactivation efficiency of bisphenols in the ZELH-zfERβ2 cell line and comparison with in human ERβ reporter gene assays from literature.

| Chemicals | ZELH-zfERβ<br>(this study) | | HELN-hERβ<br>(Grimaldi et al 2019) | | HepG2-hERβ<br>(Pelch et al 2019) | |
| --- | --- | --- | --- | --- | --- | --- |
| | EC50<br>(μM) | Maximal<br>efficacy (%) | EC50<br>(μM) | Maximal<br>efficacy (%) | EC50<br>(μM) | Maximal<br>efficacy<br>(%) |
| E2 | 0.00006 |  | 0.00009 |  | 0.0011 |  |
| BPC Cl | 0.05 | 61% | 0.046 | 35% | NT | - |
| 4-4'ODP | 0.15 | 61% | 3.06 | 72% | NT | - |
| BPC | 0.42 | 66% | 0.559 | 31% | 3.2 | 75% |
| BPF | 0.67 | 71% | 0.952 | 100% | NT | - |
| BPA | 1.36 | 55% | 0.384 | 66% | 0.35 | 97% |
| BPB | 2.57* | 66% | 0.128 | 60% | NC | 56% |
| BPS | 2.79 | 85% | 1.72 | 100% | 2.1 | 77% |
| TCBPA | 5.06 | 42% | 68 | 21% | NC | 15% |
| BPAF | 9.6* | 46% | 0.068 | 71% | 0.046 | 85% |
| BPS-MPE | NA | - | NT | - | NA | 4% |
| BPS-MAE | NA | - | NT | - | NA | 6% |

NA : non active, NT: not tested, NC: not calculated, \*: PC50 value

### The BPs are agonist ligands of zfERs and are required to induce brain aromatase

We further study the capacity of BPs to interact with the ER signalling pathway at the molecular level by performing a set of *in vivo* and *in vitro* experiments. We first performed co-exposure experiments of embryos with the ER antagonist ICI 182,780 (ICI) at 1 µM (607 µg/L) and one concentration of each bisphenol, i.e., the lowest concentration leading to maximal induction of GFP. The induction of GFP expression by BPs was partially, but not totally, downregulated in the presence of ICI (**Figure S6**) showing that functional ERs are required in mediating the induction of the brain aromatase gene by these compounds as previously demonstrated for some BPs (Le Fol et al., 2017, Pinto et al., 2019). However, by using this approach, ICI 1 μM was unable to block the effect of BPC Cl and BPS-MAE even when zebrafish embryos were pre-exposed to ICI alone during the first 48 hours of their development (data not shown). The reason of this lack of effect of ICI is not known and could simply reflect that co-exposure conditions are not optimal to efficiently block the effects especially those BPs with a high estrogenic activity such as BPC-Cl.

We also further characterized the mode of action of bisphenols on ER signalling by studying the transactivation of zebrafish ERs in the specific ZELH-zfERβ2 cell line (Cosnefroy et al., 2012). The rational for selecting this cell line was based on its (i) higher sensitivity to natural and synthetic steroidal estrogens -including 17β-estradiol, estrone, estriol and 17α ethinylestradiol (Cosnefroy et al 2012), xeno-estrogens among which BPF and BPS (LeFol et al 2017) and higher responsiveness to environmental extracts (Sonavane et al 2016) as compared to the ZELH-zfERα cell line. Furthermore, zfERβ2 is first zfER isoform to be expressed in zebrafish brain at early stages of development (Mouriec et al 2009), hence reinforcing the relevance of assessing this receptor in the present study. The results showed that, except BPS-MAE and BPS-MPE (**Figure S7B**), all the tested bisphenols induced luciferase activity in the presence of zebrafish ERs (**Figure S7A**). No induction of luciferase was observed at the test concentrations in the ZELH cell line that expresses no functional ER (data not shown), suggesting that the effects observed in ZELH-zfERβ2 cells were receptor-specific. For active BPs, EC50 values ranged from 0.05 to 9.6 µM (Table 4), which are of the same order as those reported in human *in vitro* reporter cell lines (Grimaldi et al., 2019, Pelch et al., 2019). Four out of the eleven substances had lower EC50 as compared to BPA, BPC-Cl being the most potent zfERβ2 agonist with a 26-fold higher estrogenic activity than BPA, which is consistent with previous report in human HELN ERα and ERβ cell line (Grimaldi et al., 2019). Based on EC50 values, some differences were also noticed when comparing zebrafish and human cell-based responses. For instance, BPAF appeared less active in zfERβ2 than on hERβ and hERα either in stably transfected Hela cells (Grimaldi et al., 2019) or in transiently transfected HepG2 cell (Pelch et al., 2019). Conversely, BPF and 4,4’-ODP were found much more active on zfERβ2 with 2- and 8.7-fold higher estrogenic activity as compared to BPA while they have been reported to be less active than BPA on hERβ and hERα (Grimaldi et al., 2019).

Overall, active BPs showed partial efficiency to transactivate zfERb2 reaching 42-85% of the maximum activity induced by E2 (Figure S3). This agrees with the already described partial agonist activity of BPA and bisphenol substitutes (Delfosse et al., 2012, Pinto et al., 2019, Grimaldi et al., 2019, Pelch et al., 2019). Nevertheless, some differences with human models can still be noticed since BPA, BPF or BPS were shown to act as full agonist ligand depending on the cell system used (Grimaldi et al 2019, Pelch et al., 2019, this study). Such differences in term of activity and potency depending on the cell context support the view that BPA and BPA analogues can act as selective estrogen modulators (SERM) in mammals and fish, which may account for the tissue-specific responses observed *in vivo* (Le Fol et al., 2017, Moremann et al., 2017, Pinto et al 2019, this study).

Altogether, these experiments provide further evidence that most of BPs act as agonist ligands of zebrafish ER that is required to up-regulate the *cyp1a91b* brain aromatase B expression in developing zebrafish in a concentration-dependent manner.

### Strengths and limits of the zebrafish-based embryo testing strategy

Using a large set of BPs, this work reinforces the conclusions drawn by our previous study (Christophe et al., in revision). It confirms that the combination of the two zebrafish-embryo OECD TGs (TG 236 and TG 250) using the *tg(cyp19a1b-GFP)* zebrafish line allows to inform at the same time on the toxicity, developmental effects and estrogenic activity of chemicals. By combining the two zebrafish-embryo OECD TGs (TG 236 and TG 250), a set of toxicological information was concurrently and efficiently collected (see **Table S4**) to reliably inform on the toxicity, developmental effects and estrogenic activity of selected BPs thereby providing new and relevant data, which could be used for their hazard assessment notably at the regulatory level. Together with our previous study (Christophe et al., 2022), the present work, although limited to 11 BPs, reinforces the relevance of using the *tg(cyp19a1b-GFP)* zebrafish embryo model in a unique zebrafish-embryo screening strategy (figure 4).

**Figure 4:**
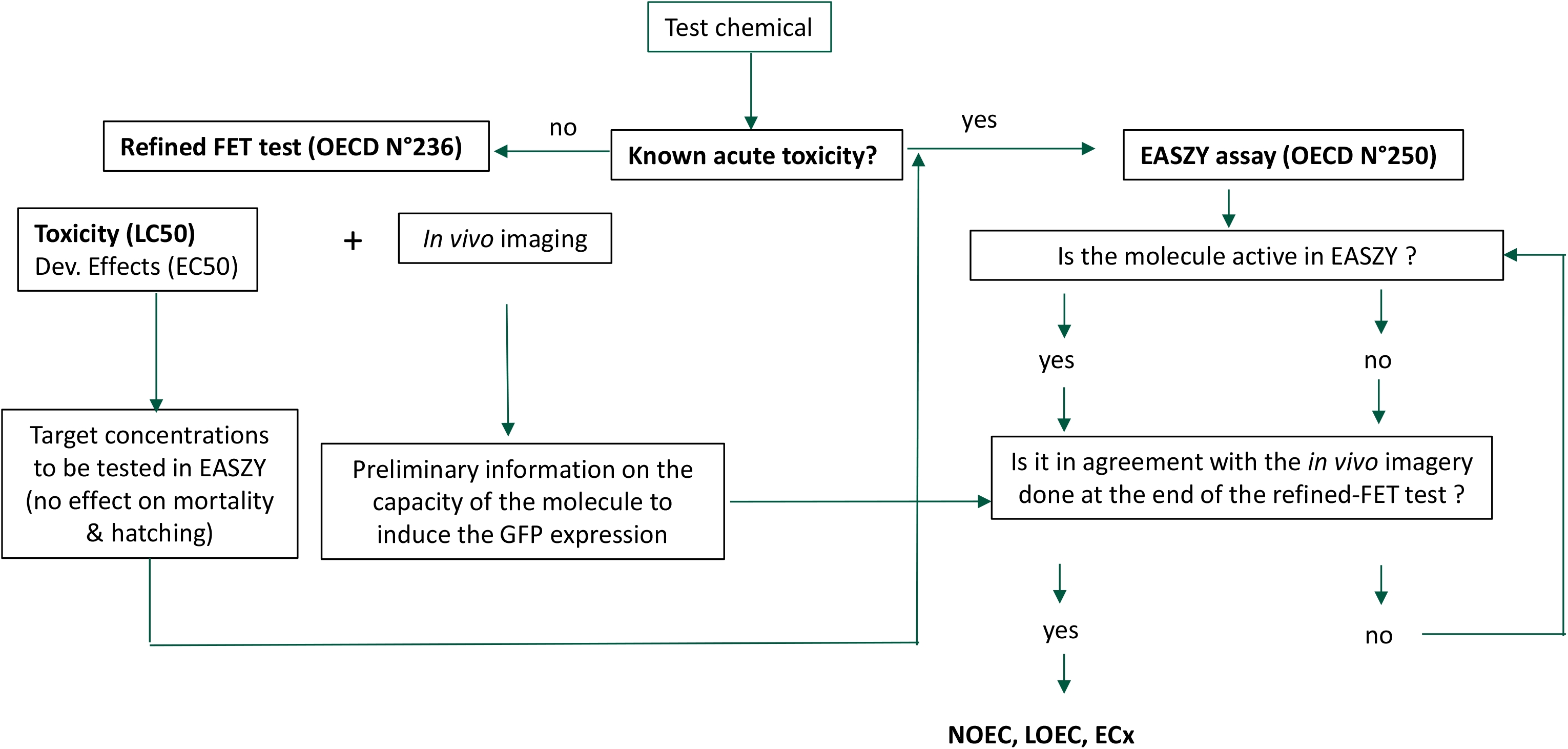
Summary of the proposed screening strategy-based on two eleuthero-embryo OECD test guidelines for the hazard assessment of chemicals

When comparing the *in vivo* and *in vitro* data, we found an overall good agreement between the *in vitro* and *in vivo* estrogenic activity both qualitatively and quantitively. It is noteworthy that chlorinated BPC, the most active bisphenol, induced *in vivo* estrogenic activity at very similar concentrations as *in vitro* (0.06 µM in EASZY and from 0.03 to 0.05 µM in cell-based assays). For the other BPs, the *in vivo* effective concentrations were 2- to 8-fold higher as compared to *in vitro* with a noticeable difference for BPS which was very much less active in embryos (Le Fol et al., 2017), reflecting the rapid *in vivo* metabolization of this compound into inactive metabolites (Le Fol et al., 2018). Interestingly, TCBPA was inactive in embryos while weakly active in zebrafish and human cell lines (this study, Grimaldi et al., 2019) but at concentrations that were toxic towards zebrafish embryos. In other words, it suggests that the environmental hazard of TCBPA relies more on its toxicity rather than on its estrogenic activity. It is also critical to emphasize that BPS-MAE and BPS-MPE were identified as inactive bisphenols in both fish and human cell-based assays (this study, Pelch et al, 2019). In contrast, BPS-MAE and BPS-MPE were clearly active *in vivo* inducing an estrogenic activity in *tg(cyp19a1:GFP)* embryos. It is now well-known that zebrafish embryos are metabolically competent having some biotransformation capacities catalyzing both Phase I and Phase II enzymatic reactions (Le Fol et al., 2018). Furthermore, previously published data have shown that EASZY assay may also be able to provide information on the estrogenic activity of pro-estrogenic chemicals, *i.e.,* that require metabolic activation prior to elicit an estrogenic response (Brion et al., 2012, Cano- Nicolau et al., 2019). Therefore, it is very likely that these two bisphenols behave as pro- estrogens, the estrogenic metabolite(s) being responsible for the ER-induction of the brain aromatase gene. To some extent, this marked difference is surprising given that the zebrafish and human hepatic cell lines have also been shown to have metabolic activities (Le Fol et al., 2015, refHepG2). Another hypothesis could rely on the fact that the estrogenic metabolite(s) produced may act as SERM as they induce an estrogenic activity in radial glial cell context but not in hepatic cells. Whatever, the differential estrogenic activity reported herein for the first time between *in vitro* ER transactivation assays and the *in vivo* EASZY assay claims for further investigation through specific metabolism and pharmacodynamic studies and highlights the importance of considering the responses in whole organisms to reliably inform about the toxicity and endocrine activity of test chemicals as done in the present work.

As previously mentioned, EASZY is the only OECD TG capable of informing on the (neuro)endocrine activity of test chemicals occurring in radial glial cells (RGCs) in a vertebrate model, but one should keep in mind that there is a gap between quantification of estrogen-like substances using EASZY and informed risk assessment. Establishing causal and possibly quantitative links between disruption of the brain aromatase gene during early brain development with adverse health effect in fish is a current challenge of research. In mammals and fish, RGCs behave as neural stem cells during embryonic development but in contrast to mammals RGCs persist in fish during the entire lifespan supporting the high neurogenic activity observed in adult fish (for review see Pellegrini *et al.,* 2016). There is some evidence showing that modulation of the brain aromatase expression or activity can impact brain proliferation and apoptosis (Diotel et al., 2013, Vaillant et al., 2021) or the behavior of exposed embryos (Kinch et al., 2018) but the underlying mechanisms mediating such events need to be further explore.

## CONCLUSION

In this study, a screening strategy based on two zebrafish eleuthero-embryos OECD test guidelines, i.e. the OECD TG 236 and TG 250, was successfully set-up and efficiently used to concurrently collect simultaneously critical data on toxicity and endocrine activity of BPs that was lacking for several of them. It allowed to produce robust data, rapidly and reliably, on the developmental effects and estrogenic activity of tested BPs, most of them being either more toxic and/or more estrogenic as BPA. Importantly, two of them (BPS-MAE and BPS-MPE) were identified as xenoestrogens while they were inactive in human and fish ER transactivation *in vitro* assays. Given the diversity of structurally or functionally bisphenol substitutes, such a screening strategy could support the current regulatory needs to assess their environmental hazard. Lastly, establishing functional and quantitative links between disruption of the brain aromatase gene and health adverse outcome(s) constitutes a major challenge with the view to potentially use EASZY data in a risk assessment perspective.

## ASSOCIATED CONTENT

### Supporting Information

Supporting Information is provided for further clarification of the experimental results. Tables describing ranges of the tested chemicals for the FET tests; ranges of the tested chemicals for the EASZY assays; summary of toxic response of fish exposed to compounds used in this study; summary table of FET and EASZY assays results. Figures showing: the modeled concentration-response curves to derive the LC50(96h); percent hatch at each time points for each chemical studied; illustration of different malformations type occurred in the FET tests; GFP fold induction above control solvent and modeling of ECx (10, 20 and 50) for BPA and BPS at the end of the FET test; GFP fold induction above control solvent and modeling of ECx (10, 20 and 50) for each chemical active on brain aromatase in EASZY assays; *in vivo* imaging of 96 hpf old live tg(*cyp19a1b*-GFP) zebrafish co-exposed to chemicals and ICI.

## AUTHOR INFORMATION

Corresponding Author *(F.B.)

## Author Contributions

All experiments of this study were designed by F.B, F.B. and S.A-A; B.P. maintained the transgenic zebrafish line in INERIS facility and produced all the zebrafish embryos in the INERIS zebrafish facility. A.C. performed the *in vivo* assays. A.C. and F.B. analyzed all data; E.M-M. ran the *in vitro* study and S.A-A. analyzed the data; A.C. draft the MS, F.B. and S.A-A. wrote the manuscript. XC, TC; XXX All authors have approved the final article.

## Notes

The authors declare no competing financial interest.

## ACKNOWLEDGMENT

This study was funded by the ANR FEATS (ANR-19-CE34-0005-05 to FB) and the MIV 26 from the French Ministry of Ecology (Characterization of endocrine disruptors to F.B., S.A. and N.H.). A.C. is funded by a doctoral research contract for INERIS (Program 190).

