## SUPPORTING INFORMATION for "A screening strategy based on two zebrafish eleuthero-embryo OECD test guidelines for the hazard assessment of chemicals: case of some bisphenol substitutes"

*Corresponding author:

**Pages: 27**

**Tables: 4**

**Figures: 6**

**Table S1.** Ranges of the tested chemicals for the FET tests.

| Chemical | Solvent | Percentage (%) | Concentrations tested (mg/L) |
|---|---|---|---|
| TCBPA | DMSO | 0.01 | 0.125, 0.25, 0.5, 1, 2 |
| BPAF | DMSO | 0.01 | 0.25, 0.5, 1, 2, 4 |
| BPS-MPE | DMSO | 0.1 | 0.62, 1.85, 5.56, 16.67, 50 |
| BPC Cl | DMSO | 0.1 | 0.99, 2.96, 8.89, 26.67, 80 |
| BPC | DMSO | 0.1 | 0.32, 0.95, 2.84, 8.53, 25.6 |
| BPB | None | - | 0.625, 1.25, 2.5, 5, 10 |
| BPA | DMSO | 0.01 | 1.25, 2.5, 5, 10, 20 |
| 4,4'ODP | DMSO | 0.01 | 0.99, 2.96, 8.89, 26.67, 80 |
| BPS-MAE | DMSO | 0.1 | 1.23, 3.7, 11.1, 33.3, 100 |
| BPF | DMSO | 0.5 | 3.75, 7.5, 15, 30, 60 |
| BPS | DMSO | 0.1 | 25, 50, 100, 200, 400 |

**Table S2**. Ranges of the tested chemicals for the EASZY assays.

| Chemical | Solvent | Percentage (%) | Concentrations tested (mg/L) |
|---|---|---|---|
| TCBPA | DMSO | 0.01 | 0.0156, 0.0312, 0.0625, 0.125, 0.25 |
| BPAF | DMSO | 0.01 | 0.021, 0.042, 0.085, 0.17, 0.34 |
| BPS-MPE | DMSO | 0.01 | 0.125, 0.25, 0.5, 1, 2 |
| BPC Cl | DMSO | 0.01 | 0.00003, 0.0003, 0.003, 0.03, 0.3 |
| BPC | DMSO | 0.01 | 0.0625, 0.125, 0.25, 0.5, 1 |
| BPB | DMSO | 0.01 | 0.078, 0.156, 0.312, 0.625, 1.25 |
| BPA | DMSO | 0.01 | 0.312, 0.625, 1.25, 2.5, 5 |
| 4,4'ODP | DMSO | 0.01 | 0.0123, 0.037, 0.11, 0.33, 1 |
| BPS-MAE | DMSO | 0.01 | 0.625, 1.25, 2.5, 5, 10 |
| BPF | DMSO | 0.01 | 0.012, 0.037, 0.11, 0.33, 1 |
| BPS | DMSO | 0.1 | 25, 50, 100, 200, 400 |

**Figure S1**. Concentration-response curves modelled using the Hill model to derive the LC50(96h) values (expressed in mg/L).

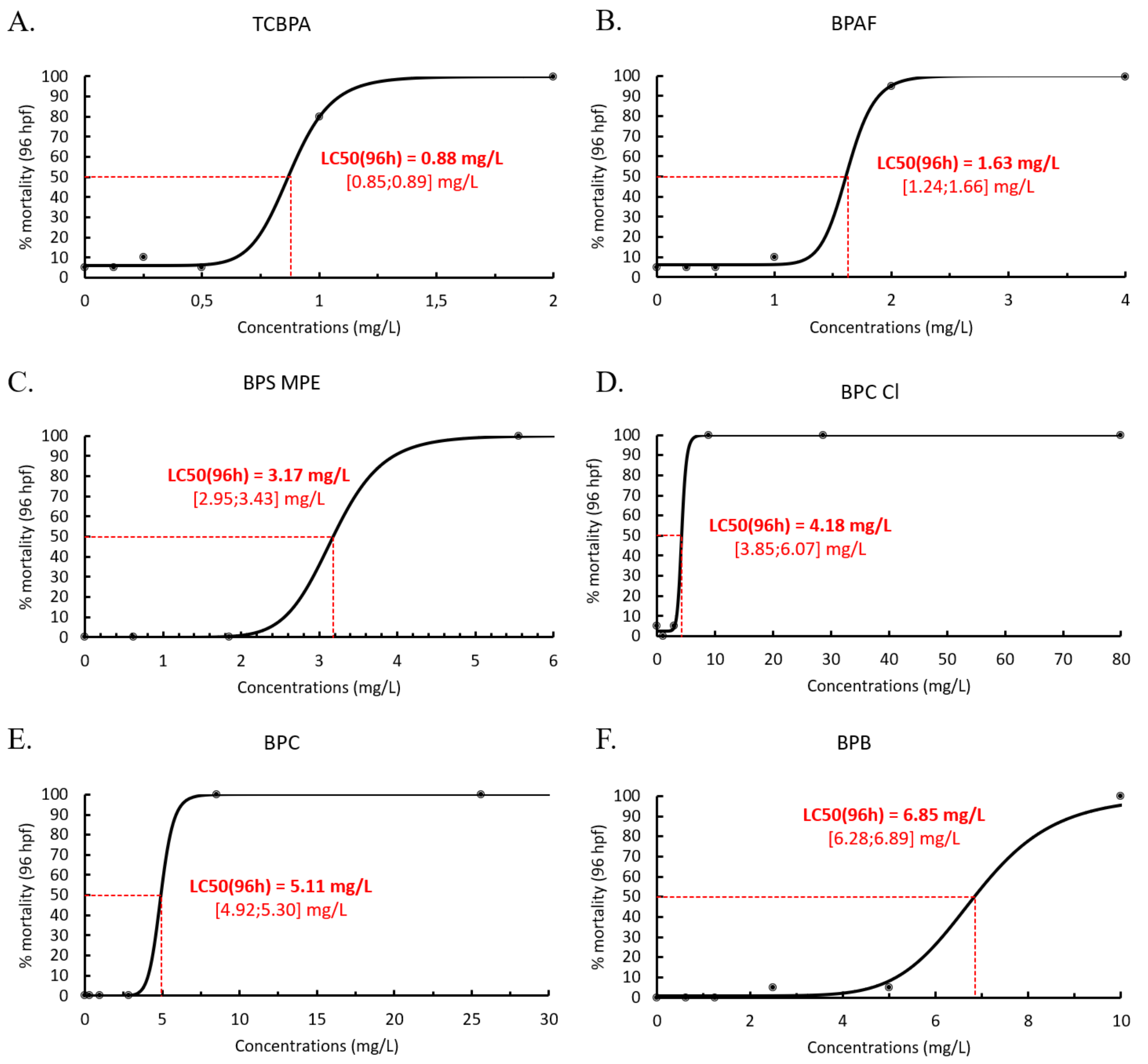

**Figure S1.** Continued

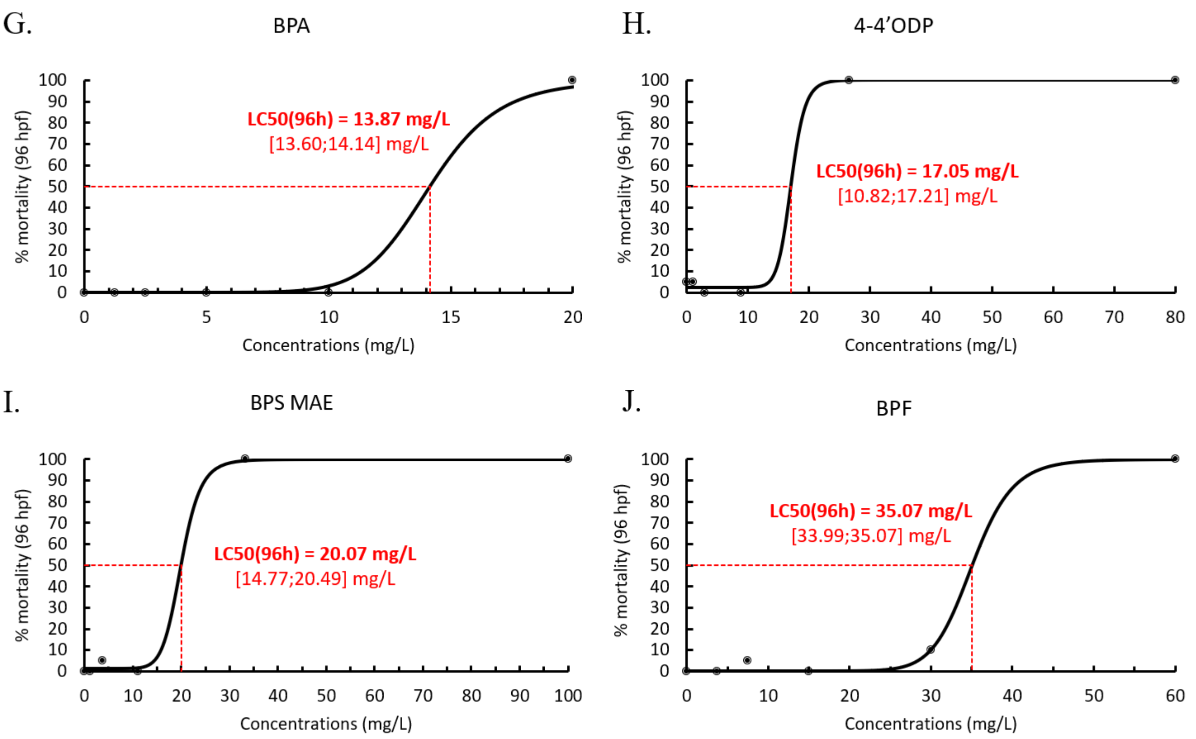

**Figure S2**. Percent hatch at each time points for each chemical studied.

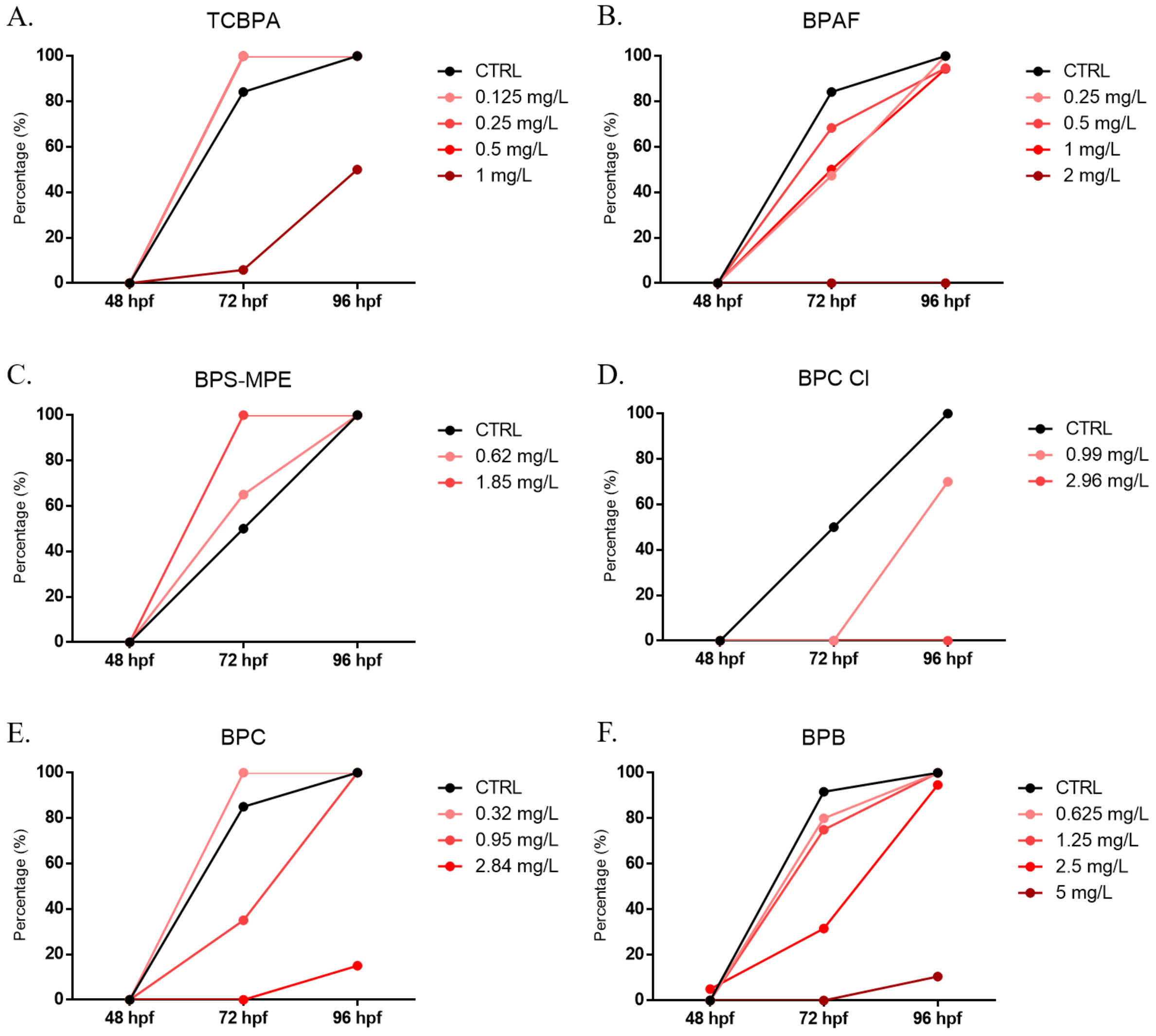

**Figure S2.** Continued

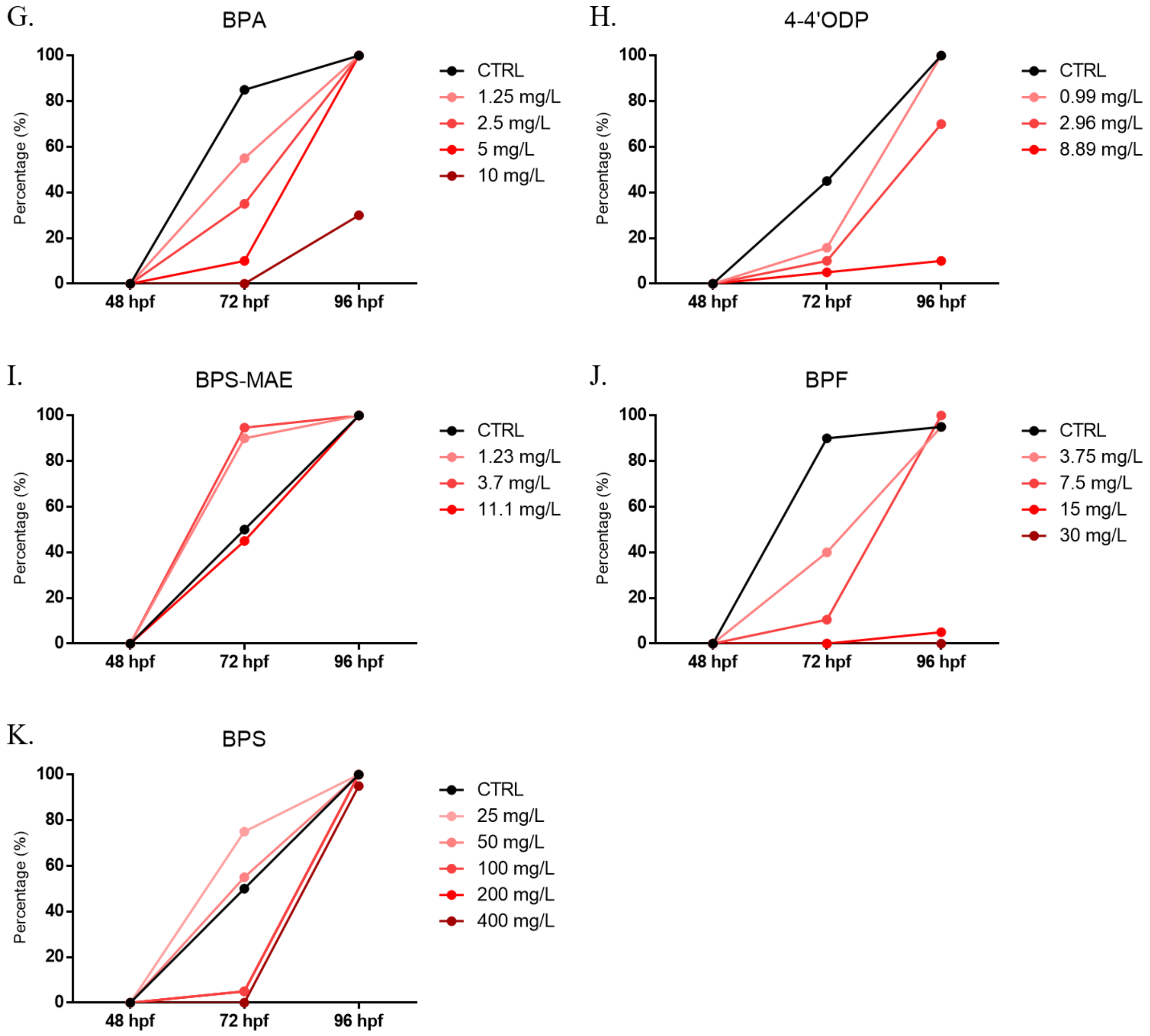

**Figure S3**. Representation of phenotypic malformations recorded in the refine FET tests. For each picture the concentration and time of exposure are indicated. deformity of yolk (*), edema (#), no or less pigmentation ($), head malformations (°), spinal malformations (¤). CTRL: control, TCBPA: tetrachlorobisphenol A, BPAF: bisphenol AF, BPB: bisphenol B, BPA: bisphenol A, 4-4'ODP: 4-4'oxydiphenol, BPF: bisphenol F. Zebrafish embryos were photographed using a X10 objective.

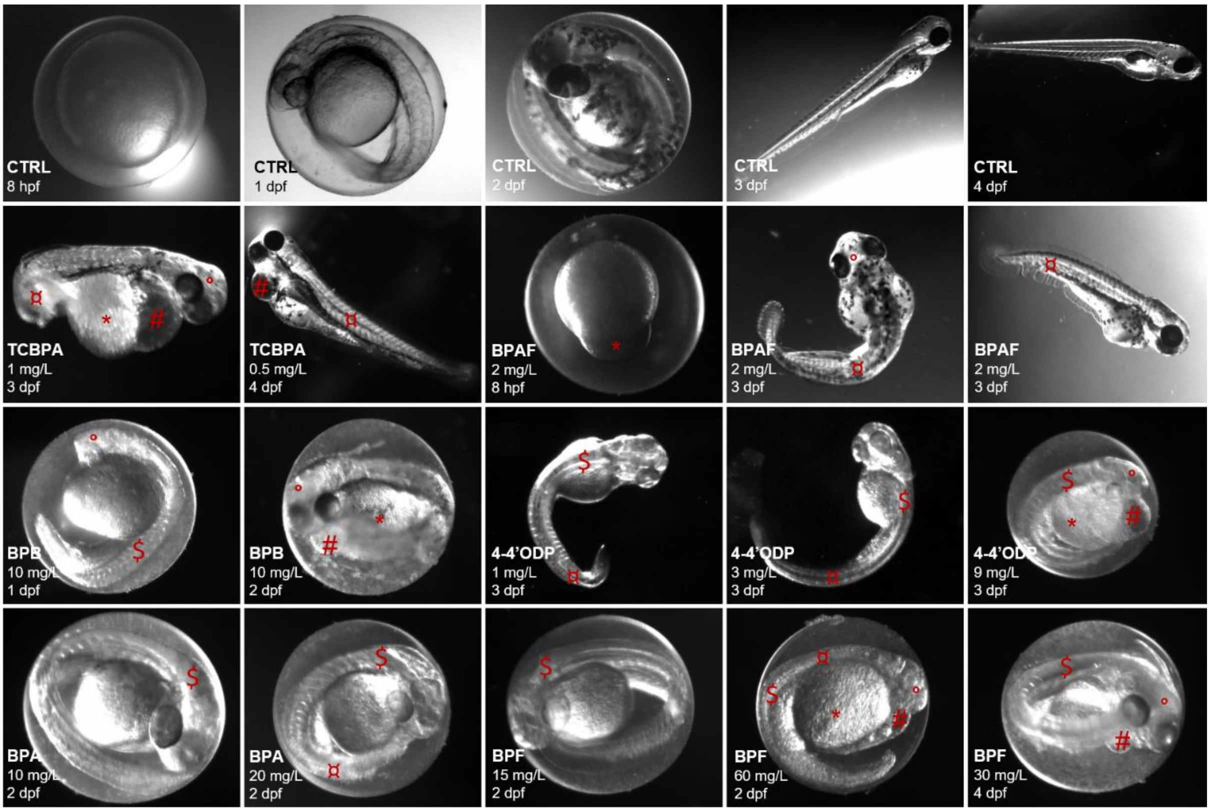

**Table S3.** Summary of the toxic responses of fish exposed to compounds used in this study. EC50 and malformations obtained in this study are reported.

| | Study | | Literature | | | |
|---|---|---|---|---|---|---|
| Chemicals | LC50(96h) [mg/L] | Malformations | Observations | Species | Test | References |
| TCBPA | 0.88 | Deformity of yolk, Spontaneous movement, Edema, Blood tail circulation, Heart rate disruption, Pigmentation, Head, Spinal malformations, delayed hatching | Concentrations above 1 μM cause lethal effects, Delayed hatching | Zebrafish Wildtype TAB and Tg(hPPARγ-eGFP) | 3-6 dpf exposure | Riu et al. 2014 |
| | | | LC50(144h) = 0.75 mg/L; LOAELs =1mg/L; Edema, Hemorrhage, Inhibition of hatching | Zebrafish embryos | 150h exposure | Song et al. 2014 |
| BPAF | 1.63 | Deformity of yolk, Spontaneous movement, Heart rate disruption, Head, Spinal malformations, Hemorrhage | No effect on survival and hatchability, Edema for 0.372 μM BPAF at 96 hpf; BPAF induced p450 aromatase, VTG and E2 at 7d | Zebrafish embryos | 7-days exposure | Chen et al. 2018 |
| | | | Malformations: Edema, Otic vesicle deformities, Delayed hatching | Zebrafish | Various (review on bisphenols) | Pelch et al. 2019 |
| | | | BPAF (0, 0.2, 0.6, 1.8, 5.2, 15.3, or 45.0 mM); NOEC = 1.8 μM; Developmental toxicity: BPAF > BPB > BPF = BPA > BPS | Zebrafish Wildtype (AB/TL) | 1-6 dpf static + 6-9 dpf semistatic | Catron et al. 2018 |
| | | | LC50(48h) = 4.2 (3.6-4.7) mg/L; Pigmentation : EC50(48h) = 3.3 (2.4-4.5) mg/L; Missing blood flow : EC50(48h) = 3.0 (0.67-13.6) mg/L; Edema : EC50(48h) = 2.8 (1.4-5.4) mg/L; Hatching inhibition: EC50(72h) = 2.2 (1.8-2.8) mg/L | Zebrafish embryos | 4-days exposure | Tisler et al. 2016 |

| | | | | | | |
|---|---|---|---|---|---|---|
| BPAF continued | | | LC50(96h) = 1.6 (0.09) mg/L; Hatching success: EC50(72h) = 0.92 (0.06) mg/L, Delayed hatching; Edema | Zebrafish embryos | 4-days exposure | Moreman et al. 2017 |
| | | | LC50(96h) = 1.95 (1.72-2.23) mg/L; Delayed hatching, Hearte rate disruption, Edema, Spinal malformations | Zebrafish embryos | 4-days exposure | Mu et al. 2018 |
| | | | LC50(144h) = 1.75 mg/L; LOAELs = 1 mg/L; Edema, Delayed hatching | Zebrafish embryos | 150h exposure | Song et al. 2014 |
| | | | LC50(48h) = 3.89 mg/L, LC50(72h) = 3.49 mg/L, LC50(96h) = 2.04 mg/L | Zebrafish embryos | 7-days exposure | Ren et al. 2017b |
| | | | Body length: NOEC = 0.5 mg/L, LOEC = 1 mg/L | Zebrafish embryos | 7-days exposure | Ren et al. 2017a |
| | | | LC50(24h) = 3.15 mg/L, LC50(48h) = 2.64 mg/L, LC50(72h) = 2.47 mg/L, LC50(96h) = 2.47 mg/L | Adult Zebrafish | 7-days exposure | Ren et al. 2017b |
| | | | Reduced body length, decreased movement distance at 120 hpf, increased number of GnRH3 neurons, increased expression of reproductive neuroendocrine-related genes and hormones: LOEC = 100 µg/L, NOEC = 1 µg/L | Zebrafish embryos | 120h-exposure | Qiu et al., 2021 |
| BPS MPE | 3.17 | Deformity of yolk, Edema, Heart rate disruption, Pigmentation, Spinal malformations, Hemorrhage | - | - | - | - |

| | | | | | | |
|---|---|---|---|---|---|---|
| BPC Cl | 4.18 | Deformity of yolk, Spontaneous movement, Edema, Blood tail circulation, Heart rate disruption, Pigmentation, Spinal malformations, Hemorrhage | - | - | - | - |
| BPC | 5.11 | Deformity of yolk, Edema, Blood tail circulation, Heart rate disruption, Spinal malformations, Hemorrhage | - | - | - | - |
| BPB | 6.85 | Deformity of yolk, Spontaneous movement, Edema, Heart rate disruption, Head malformations, Hemorrhage | Malformations: Edema, Otic vesicle deformities | Zebrafish | Various (review on bisphenols) | Pelch et al. 2019 |
| | | | BPB (0, 0.6, 1.7, 5.1, 15.0, or 44.0 mM); NOEC = 5.1 µM; Developmental toxicity: BPAF > BPB > BPF = BPA > BPS | Zebrafish Wildtype (AB/TL) | 1-6 dpf static + 6-9 dpf semistatic | Catron et al. 2018 |
| | | | Yeast two-hybrid assay, estrogenicity assessment: BPB ≥ BPA, BPF > BPS; decrease of activity probably due to higher acute toxicity | Daphnia magna | acute toxicity assay | Chen et al. 2001 |
| | | | LC50(24h) = 7.18 mg/L, LC50(48h) = 6.54 mg/L, LC50(72h) = 4.53 mg/L, LC50(96h) = 3.88 mg/L | Zebrafish embryos | 7-days exposure | Ren et al. 2017b |
| | | | Body length: NOEC = 1 mg/L, LOEC = 2 mg/L | Zebrafish embryos | 7-days exposure | Ren et al. 2017a |
| | | | LC50(24h) = 5.07 mg/L, LC50(48h) = 4.64 mg/L, LC50(72h) = 4.15 mg/L, LC50(96h) = 4.15 mg/L | Adult Zebrafish | 7-days exposure | Ren et al. 2017b |

| | | | | | | |
|---|---|---|---|---|---|---|
| BPB continued | | | Reduced body length, decreased movement distance at 120 hpf, increased number of GnRH3 neurons, increased expression of reproductive neuroendocrine-related genes and hormones: LOEC = 100 µg/L, NOEC = 1 µg/L | Zebrafish embryos | 120h-exposure | Qiu et al., 2021 |
| BPA | 13.87 | Deformity of yolk, Growth retardation, Spontaneous movement, Edema, Heart rate disruption, Pigmentation, Spinal malformations, Hemorrhage | 100 µM (23 mg/L): 100% mortality at 72h, developmental and comportemental effects; 100 nM (0,023 mg/L): no effect of BPA | Zebrafish embryos | 5-days exposure | Björnsdotter et al. 2017 |
| | | | LC25 = 2.8x10-6 M, NOAEL ≥ 10-6 M, non-teratogen | Zebrafish embryos | 120h exposure | Brannen et al. 2010 |
| | | | EC50 = 1226 µg/L [1169;1310] = 1370 nM, LOEC = 1000 µg/L = 4380.32 nM, No developmental toxicity | Zebrafish embryos ERE-GFP-Casper | 120h exposure | Green et al. 2016 |
| | | | LC50(24h) = 16.75 mg/L, NOEC = 2 mg/L, 100% mortality at 25 mg/L; Edema, Blood flow, Delayed hatching | Zebrafish embryos | 72h exposure | Duan et al. 2008 |
| | | | Exposure to 20 mg/L of BPA; 24h: Deformity of yolk, Edema, Growth retardation, Head, Spinal malformations; 48h: 65% mortality, Heart rate disruption, Head malformations, Pigmentation, Edema | Zebrafish Wildtype (AB/TL) | 48h exposure | Makarova et al. 2016 |

| | | | | |
|---|---|---|---|---|
| BPA continued | LC50(24h) = 9.503 mg/L [9.184;10.565]; LC50(48h) = 8.688 mg/L [8.560;8.811]; LC50(72h) = 7.681 mg/L [7.414;7.989]; LC50(96h) = 6.669 [5.654;7.607]; Edema: 100% at 96h for 8 and 9 mg/L | Zebrafish larvae (3 dpf) | 4-days exposure | Thi et al. 2016 |
| | 235 mg/L: 100% mortality at 24h; 135 mg/L: 100% mortality at 48h; 50 mg/L: malformations at 24h, no heart beat at 48h, Hemorrhage, Pigmentation, 100% mortality at 72h; 16 and 14 mg/L: Pigmentation, Hemorrhage, Edema, Heart rate disruption at 48h, 80% mortality at 72h; 2 mg/L: no effect | Zebrafish embryos | 8-days exposure | Ortiz et al. 2009 |
| | No effect of BPA for hatchability, phenotype and mortality with exposure at 24-48 hpf and 0.4 µM | Zebrafish embryos | 24-48h exposure | Huang et al. 2016 |
| | LC50(96h) = 12.8 mg/L | Zebrafish | 4-days exposure | Corrales et al. 2016 |
| | LC50(96h) = 4.2 mg/L | Fathead minnow | 4-days exposure | Corrales et al. 2016 |
| | LC50 = 17.5 ± 0.37 µM (= 3.995 ± 0.08 mg/L); 5-15 µM induced 4-8% mortality; Edema, Hemorrhage, Spinal malformations, Delayed hatching | Zebrafish embryos | 5-days exposure | McCormick et al. 2010 |

| | | | | |
|---|---|---|---|---|
| BPA continued | LC50(5dpf) = 5 (0.89) mg/L, LC50(28dpf) = 1.8 (0.23) mg/L, LOAEL(5dpf) = 1 mg/L, LOAEL(28dpf) = nd; Edema, Hemorrhage, Curved tails, Delayed hatching | Zebrafish embryos | 7-days exposure | McCormick et al. 2011 |
| | 1000 µg/L induce no mortality at 120 hpf, increase hatching at 48 and 54 hpf, Body length reduced in concentration-dependent manner | Zebrafish embryos | 120h exposure | Qiu et al. 2018 |
| | BPA (1, 5, and 15 µM) have no effect on the development between 8 and 96 hpf; BPA alters motor behavior of larvae between 18 and 120 hpf | Zebrafish embryos (Wildtype AB) | 120h exposure | Wang et al. 2013 |
| | Exposure to 10-100 µM of BPA; No mortality until 70 µM, no malformations below 30 µM; 30-70 µM : Edema, Craniofacial abnormalities, Delayed hatching; Transient effect on larval hyperactivity and learning to select correct arm in T-maze to avoid an electric shock for 0.01 and 0.1 µM (no effect for 1 and 10 µM) | Zebrafish embryos | 120h exposure | Saili et al. 2012 |
| | Exposure to 0.1, 1, 10, 100, 1000 µg/L with renewal every 12h and final DMSO concentration at 0.005% v/v; No effect on survival and time of hatching; Malformations: Spinal curvature, Edema | Zebrafish embryos (Wildtype AB) | 4-168h exposure | Wu et al. 2011 |

| | | | | |
|---|---|---|---|---|
| BPA continued | BPA induced GFP in liver and heart; BPA has effects on the heart, in its structure but not in its cardiovascular function | Zebrafish embryos/larvae TG(ERE:GFP) | 5-days exposure or 6-15 dpf exposure | Brown et al. 2019 |
| | Malformations: Edema, Otic vesicle deformities | Zebrafish | Various (review on bisphenols) | Pelch et al. 2019 |
| | BPA (0, 0.2, 0.6, 1.7, 2.9, 5.7, 11.5, 23.0, or 45.0 mM); NOEC = 11.5 µM; Developmental toxicity: BPAF > BPB > BPF = BPA > BPS | Zebrafish Wildtype (AB/TL) | 1-6 dpf static exposure + 6-9 dpf semistatic exposure | Catron et al. 2018 |
| | Yeast two-hybrid assay, estrogenicity assessement: BPB ≥ BPA, BPF > BPS | Daphnia magna | acute toxicity assay | Chen et al. 2001 |
| | Renewal of test media every 12h, higher hatching rates at 48 and 55 hpf for 1 and 10 µg/L, no effect on survival until 1000 µg/L | Zebrafish GnRH3-EMD transgenic | 120h exposure | Qiu et al. 2016 |
| | LC50(48h) = 15.9 (13.8-18.3) mg/L; Pigmentation : EC50(48h) = 3.6 (2.5-5.3) mg/L; Missing blood flow : EC50(48h) = 12.3 (10.3-14.6) mg/L; Edema : EC50(48h) = 13.9 (8.5-22.8) mg/L; Hatching inhibition: EC50(72h) = 4.0 (3.1-5.2) mg/L | Zebrafish embryos | 4-days exposure | Tisler et al. 2016 |
| | LC50(96h) = 10.4 (9.44-11.58) mg/L; Edema | Zebrafish embryos | 4-days exposure | Mu et al. 2018 |
| | LC50(144h) = 7.5 mg/L; LOAELs = 5 mg/L | Zebrafish embryos | 150h exposure | Song et al. 2014 |

| | | | | |
|---|---|---|---|---|
| BPA continued | LC50(96h) = 12 (0.22) mg/L; Hatching success: EC50(72h) = 5;7 (0.33) mg/L, Delayed hatching; Edema, Craniofacial abnormality, Hemorrhage | Zebrafish embryos | 4-days exposure | Moreman et al. 2017 |
| | LC50(96h) = 9.82 mg/L, LC10(96h) = 6.62 mg/L | Zebrafish embryos | 4-days exposure | Blanc et al. 2019 |
| | LC50(96h) = 35 µM = 8.041 mg/L [7.846;8.24] mg/L, delayed hatching: EC50(96h) = 5.25 mg/L [2.982;5.819] mg/L | Zebrafish embryos | 4-days exposure | Chow et al. 2013 |
| | Mortality at 219 µM (50 mg/L), delayed hatching from 35 µM (8.041 mg/L), inflated swim bladder from 17.5 µM (4 mg/L), scoliosis (16% at 8.041 mg/L); Reduced body length, deformity of yolk, head and spinal malformations, pigmentation (malformations occurring at low concentrations) | Zebrafish embryos | 2-5 dpf exposure | Martinez et al. 2019 |
| | Body length: NOEC = 2 mg/L, LOEC = 4 mg/L | Zebrafish embryos | 7-days exposure | Ren et al. 2017a |
| | LC50(24h) = 9.51 mg/L, LC50(48h) = 9.31 mg/L, LC50(72h) = 8.09 mg/L, LC50(96h) = 8.09 mg/L | Adult Zebrafish | 7-days exposure | Ren et al. 2017b |
| | Malformations observed: axial malformation, pericardial edema, yolk sac edema (72 hpf); exposure to 0.001 mg/L of BPA | Zebrafish embryos | 72h-exposure | Ustundag et al. 2017 |

| | | | | | | |
|---|---|---|---|---|---|---|
| BPA continued | | | Accelerated hatching, reduced body length, decreased movement distance at 120 hpf, increased number of GnRH3 neurons, increased expression of reproductive neuroendocrine-related genes and hormones: LOEC = 100 μg/L, NOEC = 1 μg/L | Zebrafish embryos | 120h-exposure | Qiu et al., 2021 |
| 4-4'ODP | 17.05 | Deformity of yolk, Growth retardation, Edema, Blood tail circulation, Heart rate disruption, Pigmentation, Head, Spinal malformations, Hemorrhage | - | - | - | - |
| BPS MAE | 20.07 | Deformity of yolk, Spontaneous movement, Edema, Blood tail circulation, Hearte rate disruption, Pigmentation, Spinal malformations | - | - | - | - |
| BPF | 35.07 | Deformity of yolk, Spontaneous movement, Edema, Blood tail circulation, Heart rate disruption, Pigmentation, Head, Spinal malformations, Hemorrhage | 1000 μg/L: no mortality at 120 hpf, increase hatching at 48 and 54 hpf; body length reduced in concentration-dependent manner | Zebrafish embryos | 120h-exposure | Qiu et al. 2018 |
| | | | LC50(96h) = 10.030 μg/L; Delayed hatching, Edema, Spinal malformations, Hemorrhage, Coagulation | Zebrafish embryos (Wildtype AB) | 120h-exposure | Yang et al. 2018 |

| | | | | |
|---|---|---|---|---|
| BPF continued | Developmental effects: Pigmentation, Heart rate disruption, Spontaneous movement, Delayed hatching, Spinal malformations | Zebrafish embryos | 96h-exposure | Mu et al. 2019 |
| | BPF (0, 0.2, 0.6, 1.8, 5.2, 15.3, 45.0 mM); NOEC = 15.3 µM; Developmental toxicity: BPAF > BPB > BPF = BPA > BPS | Zebrafish Wildtype (AB/TL) | 1-6 dpf + 6-9 dpf | Catron et al. 2018 |
| | Only 20% mortality at 20 mg/L; Pigmentation: EC50(48h) = 1.1 (0.92-1.3) mg/L; Edema: EC50(48h) = 10.7 (9.4-12.2) mg/L; Hatching inhibition: EC50(72h) = 6.8 (5.8-8.5) mg/L | Zebrafish embryos | 4-days exposure | Tisler et al. 2016 |
| | LC50(96h) = 32 (0.55) mg/L; Hatching success: EC50(72h) = 14 (0.41) mg/L, Delayed hatching; Edema, Craniofacial abnormality, Tail development, Hemorrhage, Deformity of yolk | Zebrafish embryos | 4-days exposure | Moreman et al. 2017 |
| | LC50(96h) = 19.6 (18.47-20.67) mg/L; Delayed hatching, Hearte rate disruption, Edema, Spinal malformations, Pigmentation | Zebrafish embryos | 4-days exposure | Mu et al. 2018 |
| | LC50(24h) = 9.13 mg/L, LC50(48h) = 8.93 mg/L, LC50(72h) = 8.56 mg/L, LC50(96h) = 7.40 mg/L | Zebrafish embryos | 7-days exposure | Ren et al. 2017b |
| | Body length: NOEC = 6 mg/L, LOEC = 8 mg/L | Zebrafish embryos | 7-days exposure | Ren et al. 2017a |

| | | | | | | |
|---|---|---|---|---|---|---|
| BPF continued | | | LC50(24h) = 10.10 mg/L, LC50(48h) = 9.86 mg/L, LC50(72h) = 9.51 mg/L, LC50(96h) = 9.51 mg/L | Adult Zebrafish | 7-days exposure | Ren et al. 2017b |
| | | | LC50(96h) = 7.40 mg/L | | | Gu et al. (2020) |
| | | | Accelerated hatching, reduced body length, increased number of GnRH3 neurons, increased expression of reproductive neuroendocrine-related genes and hormones: LOEC = 100 µg/L, NOEC = 1 µg/L | Zebrafish embryos | 120h-exposure | Qiu et al., 2021 |
| BPS | CNC | Deformity of yolk | 1000 µg/L: no mortality at 120 hpf, increase hatching at 48 and 54 hpf, Body length reduced in concentration-dependent manner | Zebrafish embryo | 120h exposure | Qiu et al. 2018 |
| | | | Malformations: Edema, Otic vesicle deformities, Delayed hatching | Zebrafish | Various (review on bisphenols) | Pelch et al. 2019 |
| | | | BPS (0, 0.2, 0.6, 1.8, 5.2, 15.3, 45.0 mM); NOEC = 45 µM, no toxicity for BPS; Developmental toxicity: BPAF > BPB > BPF = BPA > BPS | Zebrafish Wildtype (AB/TL) | 1-6 dpf static exposure + 6-9 dpf semi-static exposure | Catron et al. 2018 |
| | | | Yeast two-hybrid assay, estrogenicity assessement: BPB ≥ BPA, BPF > BPS | Daphnia magna | acute toxicity assay | Chen et al. 2001 |
| | | | BPS (0, 1, 3, 10, 30 µg/L), DMSO 0.01% (v/v); No effect on survival or growth at 168 hpf, no significative malformations, Delayed hatching | Zebrafish embryos (Wildtype AB) | 168h exposure | Zhang et al. 2017 |

| | | | | |
|---|---|---|---|---|
| BPS continued | Renewal of test media every 12h, no mortality for 100 µg/L | Zebrafish GnRH3-EMD transgenic | 120h exposure | Qiu et al. 2016 |
| | LC50(96h) = 199 (7.6) mg/L; Hatching success: EC50(72h) = 155 (15) mg/L, Delayed hatching; Edema, Craniofacial abnormality, Tail development | Zebrafish embryos | 4-days exposure | Moreman et al. 2017 |
| | No mortality and no malformations until 50 mg/L | Zebrafish embryos | 4-days exposure | Mu et al. 2018 |
| | LC50(96h) > 400 µM (> 100 mg/L), LC10(96h) > 400 µM (> 100 mg/L) | Zebrafish embryos | 4-days exposure | Blanc et al. 2019 |
| | No mortality, LC50(96h) > 30 µM (> 7.5 mg/L) | Zebrafish embryos | 4-days exposure | Le Fol et al. 2017b |
| | LC50(24h) = 361 mg/L, LC50(48h) = 346 mg/L, LC50(72h) = 331 mg/L, LC50(96h) = 323 mg/L | Zebrafish embryos | 7-days exposure | Ren et al. 2017b |
| | Heart rate: EC50(78h) = 318 mg/L; Hatching rate: EC50(120h) = 200 mg/L; Body length: NOEC = 300 mg/L, LOEC = 350 mg/L | Zebrafish embryos | 7-days exposure | Ren et al. 2017a |
| | LC50(24h) = 343 mg/L, LC50(48h) = 343 mg/L, LC50(72h) = 343 mg/L, LC50(96h) = 343 mg/L | Adult Zebrafish | 7-days exposure | Ren et al. 2017b |
| | Accelerated hatching, reduced body length, increased number of GnRH3 neurons, increased expression of reproductive neuroendocrine-related genes and hormones: LOEC = 100 µg/L, NOEC = 1 µg/L | Zebrafish embryos | 120h-exposure | Qiu et al., 2021 |

42

**Figure S4.** GFP fold induction above control solvent and modelization of the ECx (10, 20 and 50) for BPA and BPS at the end of the FET test.

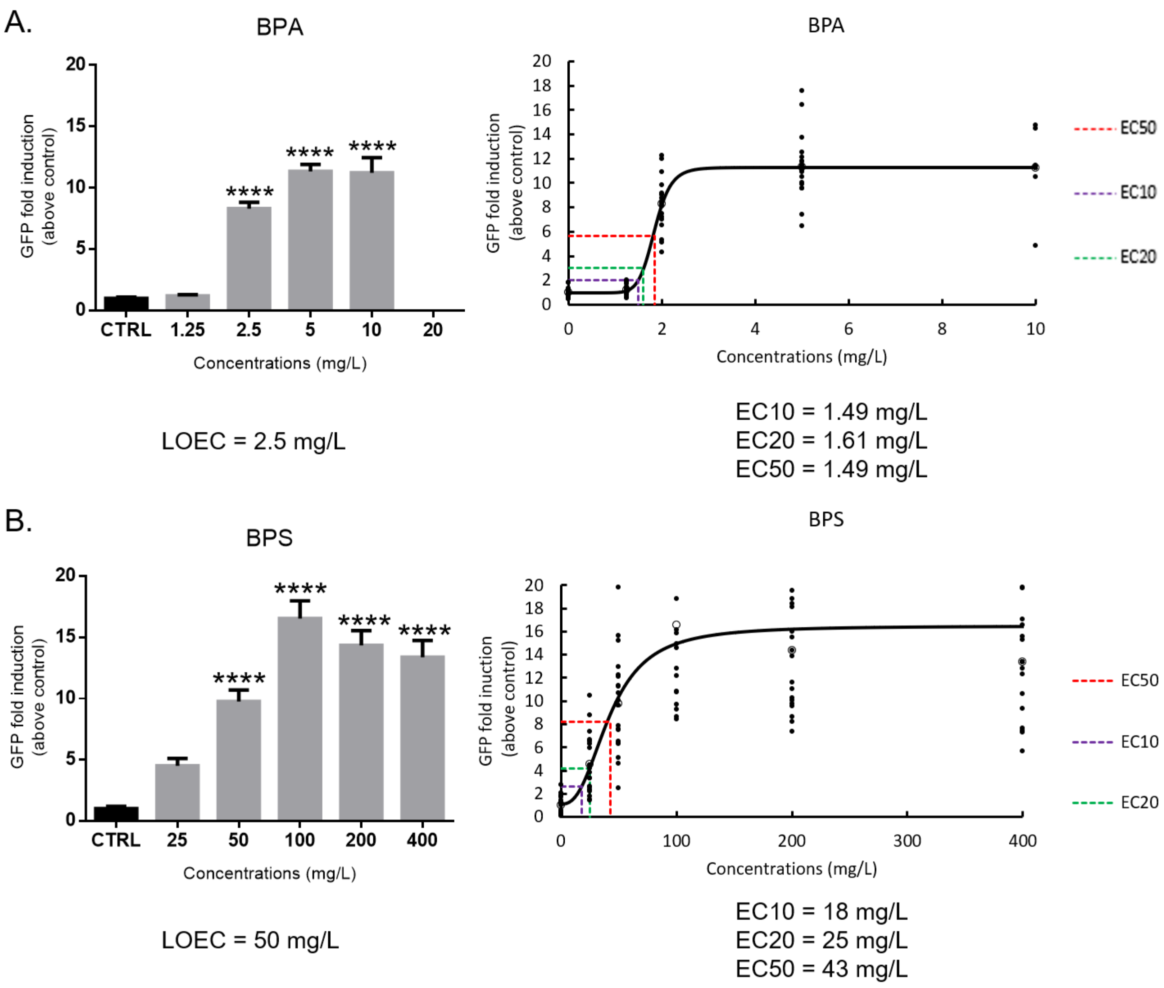

**Figure S5**. GFP fold induction above control solvent and modelization of the ECx (10, 20 and 50) for each chemical active on brain aromatase in the EASZY assay.

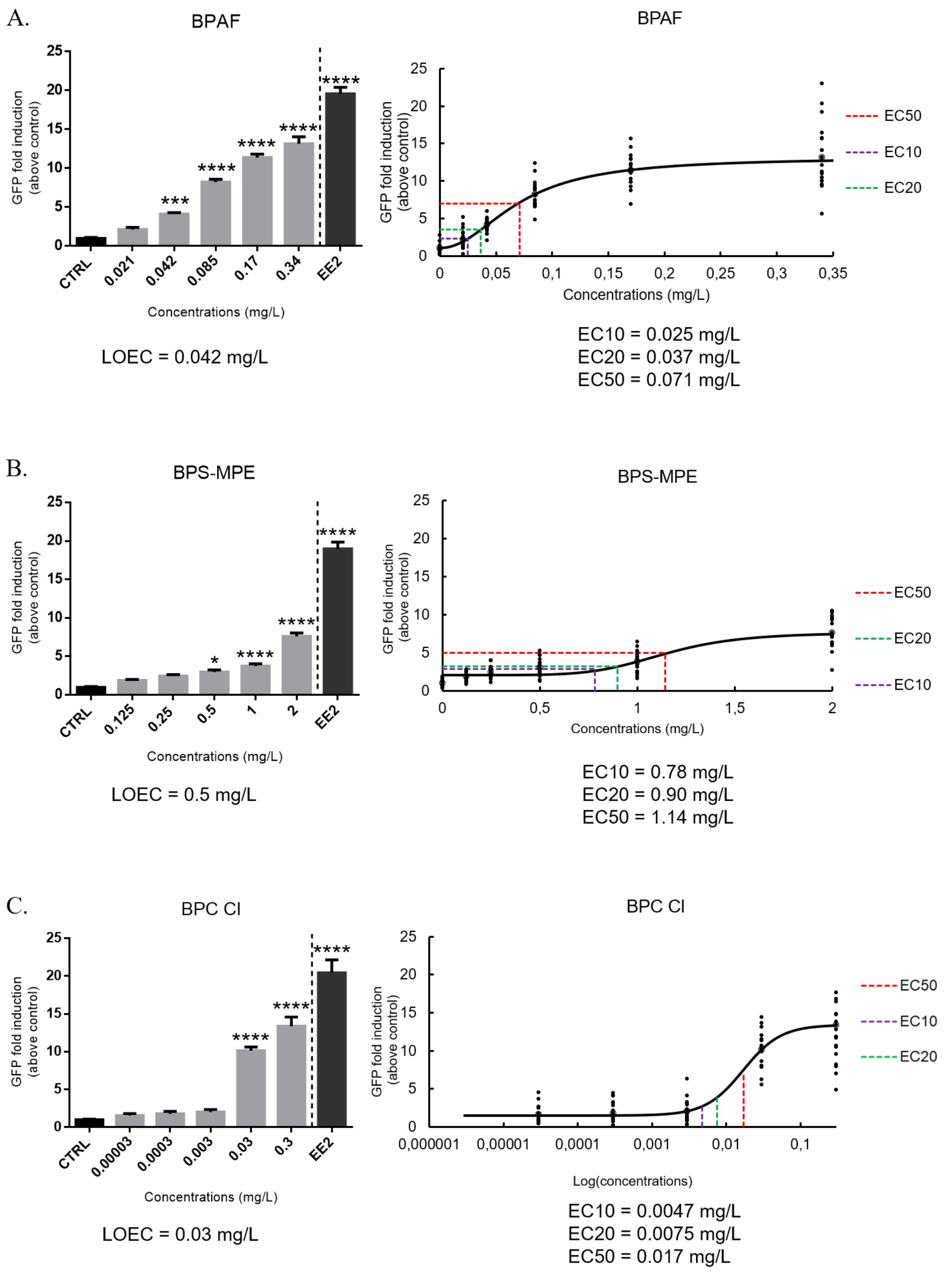

**Figure S5.** Continued

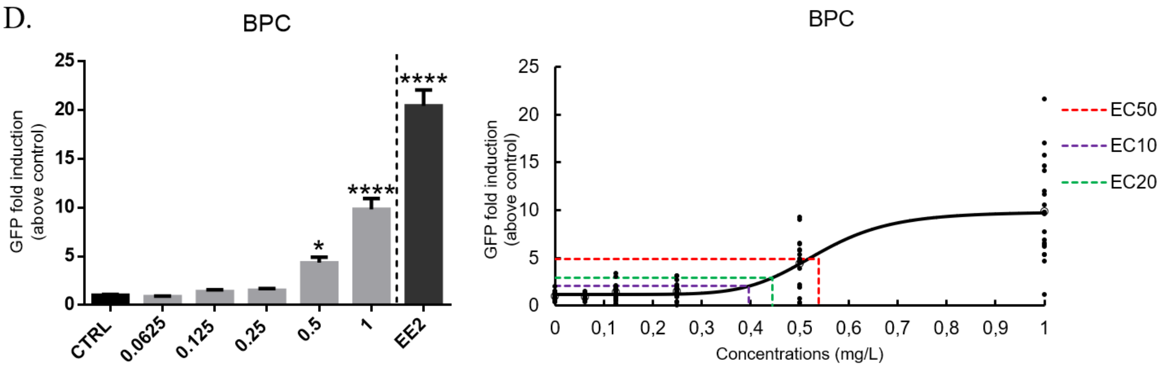


LOEC = 0.5 mg/L

EC10 = 0.40 mg/L
EC20 = 0.44 mg/L
EC50 = 0.54 mg/L

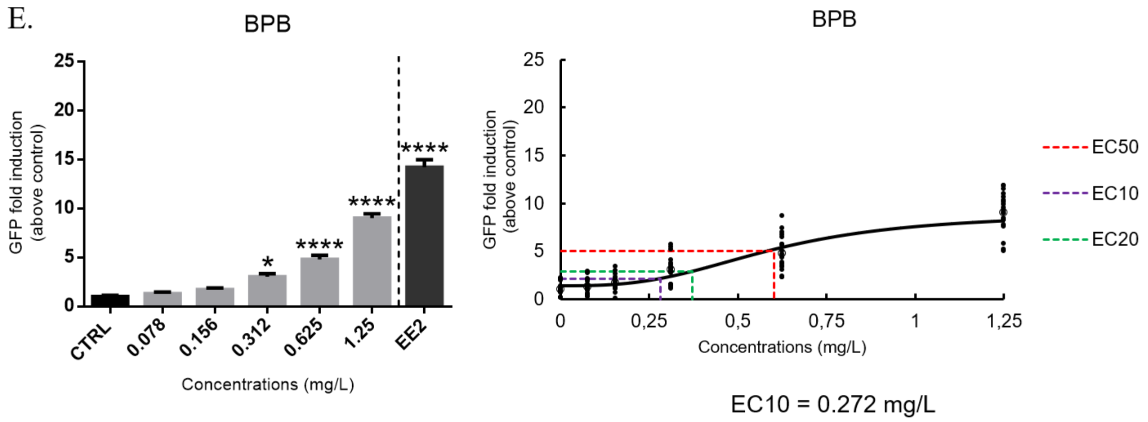


LOEC = 0.312 mg/L

EC10 = 0.272 mg/L
EC20 = 0.363 mg/L
EC50 = 0.596 mg/L

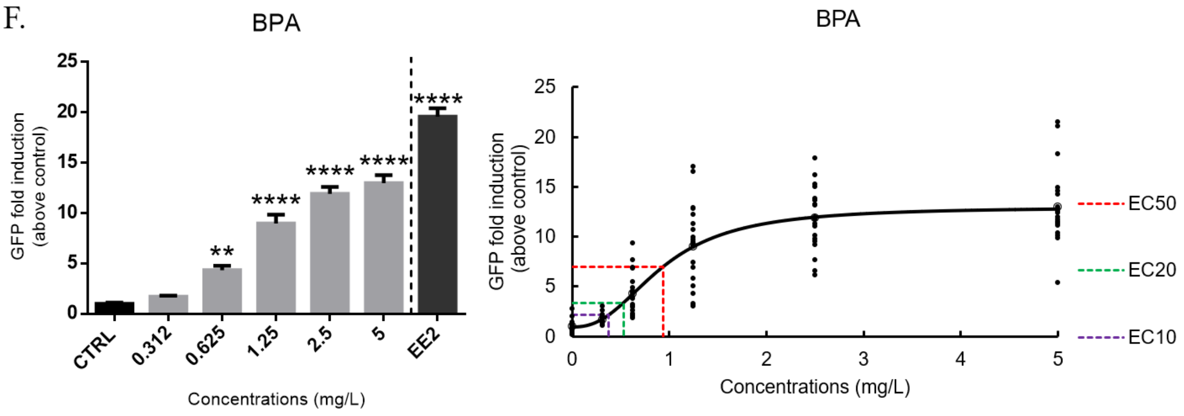


LOEC = 0.625 mg/L

EC10 = 0.38 mg/L
EC20 = 0.53 mg/L
EC50 = 0.93 mg/L

**Figure S5.** Continued

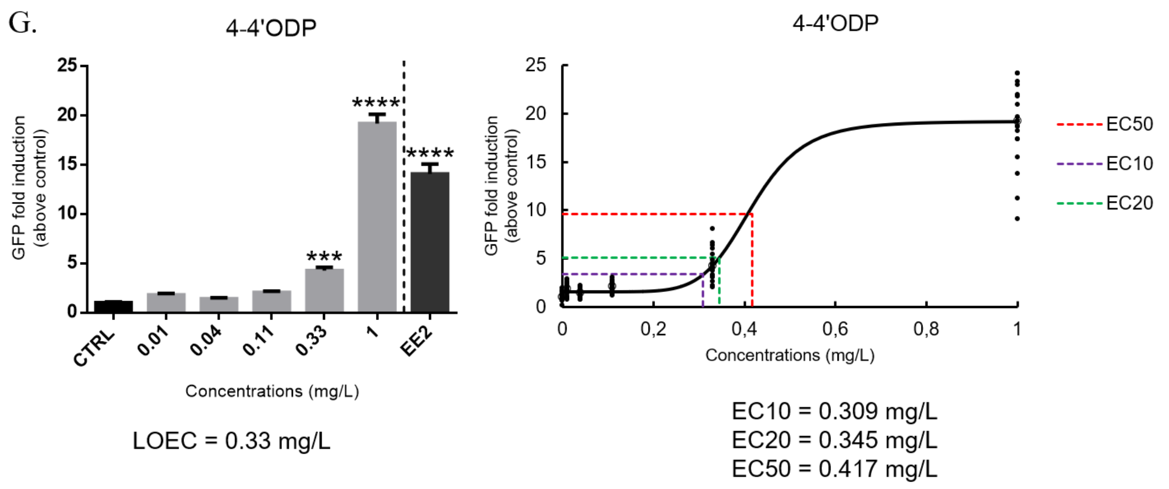


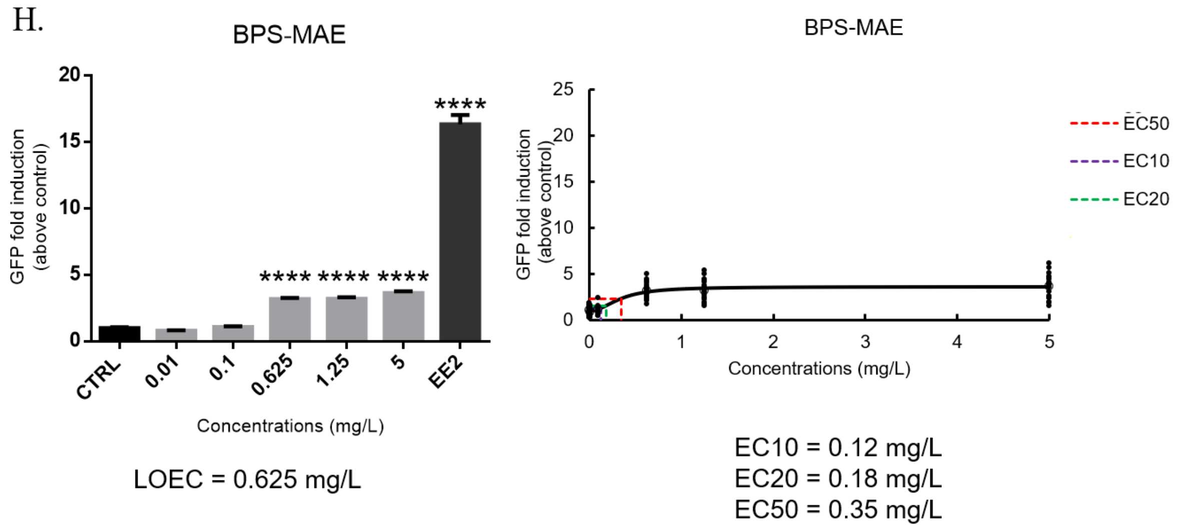


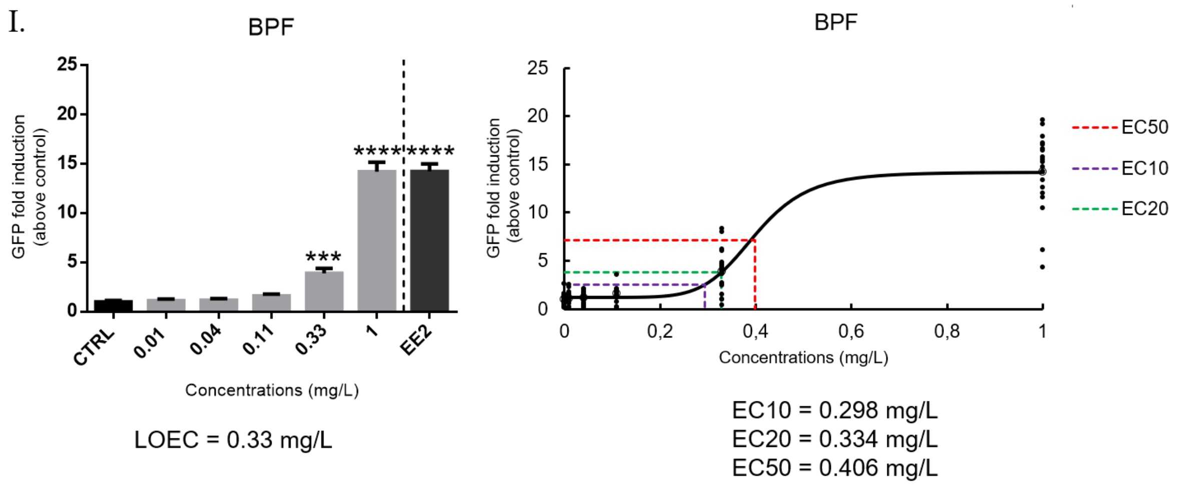

**Figure S5.** Continued

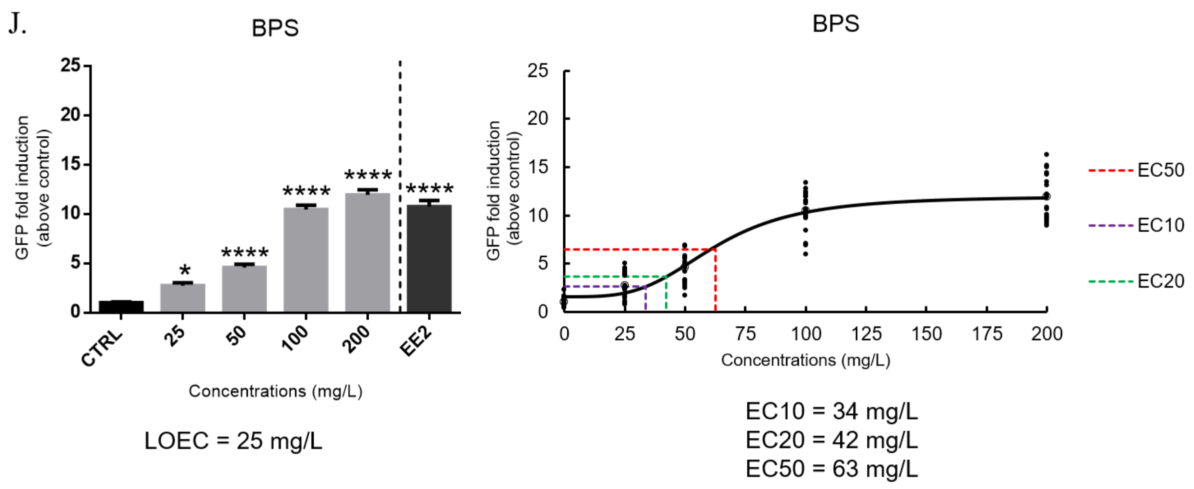

**Table S4**. Summary table of FET and EASZY assays results. **For the FET test**, LC50(96h) are reported as the smallest LOECs obtained for malformations and potential estrogenic activity detected at the end of the test with fluorescence imagery at 96h (-: inactive, +: active). For BPS, a complete concentration-response was obtained at the end of the FET test and EC50 can be modelized (Figure S3). **For the EASZY assay**, the lowest observed effect concentrations (LOECs) inducing a significant GFP induction above control are reported as well as the effective concentrations leading to 50% of effect (EC50 expressed in mg/L and nM).

| | FET assay (OECD 236) | | | EASZY assay (OECD 250) | | |
|---|---|---|---|---|---|---|
| Chemicals | Acute toxicity LC50 (mg/L) | Developmental effects LOECs (mg/L) | GFP induction (estrogenicity) | LOECs (mg/L) | EC50 (mg/L) | EC50 (nM) |
| TCBPA | 0.88 | 0.5 | - | n.a. | n.a. | n.a. |
| BPAF | 1.63 | 1 | + | 0.042 | 0.071 | 211 |
| BPS-MPE | 3.17 | 5.56 | + | 0.5 | 1.14 | 3.35 |
| BPC Cl | 4.18 | 0.99 | + | 0.03 | 0.017 | 0.0604 |
| BPC | 5.11 | 2.84 | + | 0.5 | 0.54 | 2.11 |
| BPB | 6.85 | 5 | + | 0.312 | 0.596 | 2460 |
| BPA | 13.87 | 10 | EC50 = 1.49 mg/L | 0.625 | 0.93 | 4063 |
| 4-4'ODP | 17.05 | 0.99 | + | 0.33 | 0.417 | 2062 |
| BPS-MAE | 20.07 | 11.1 | + | 0.625 | 0.35 | 1205 |
| BPF | 35.07 | 3.75 | + | 0.33 | 0.406 | 2030 |
| BPS | > 100 | >100 | EC50 = 43 mg/L | 25 | 63 | 249326 |

**Figure S6**. Imaging results of co-exposition with bisphenols and ICI 182,780.

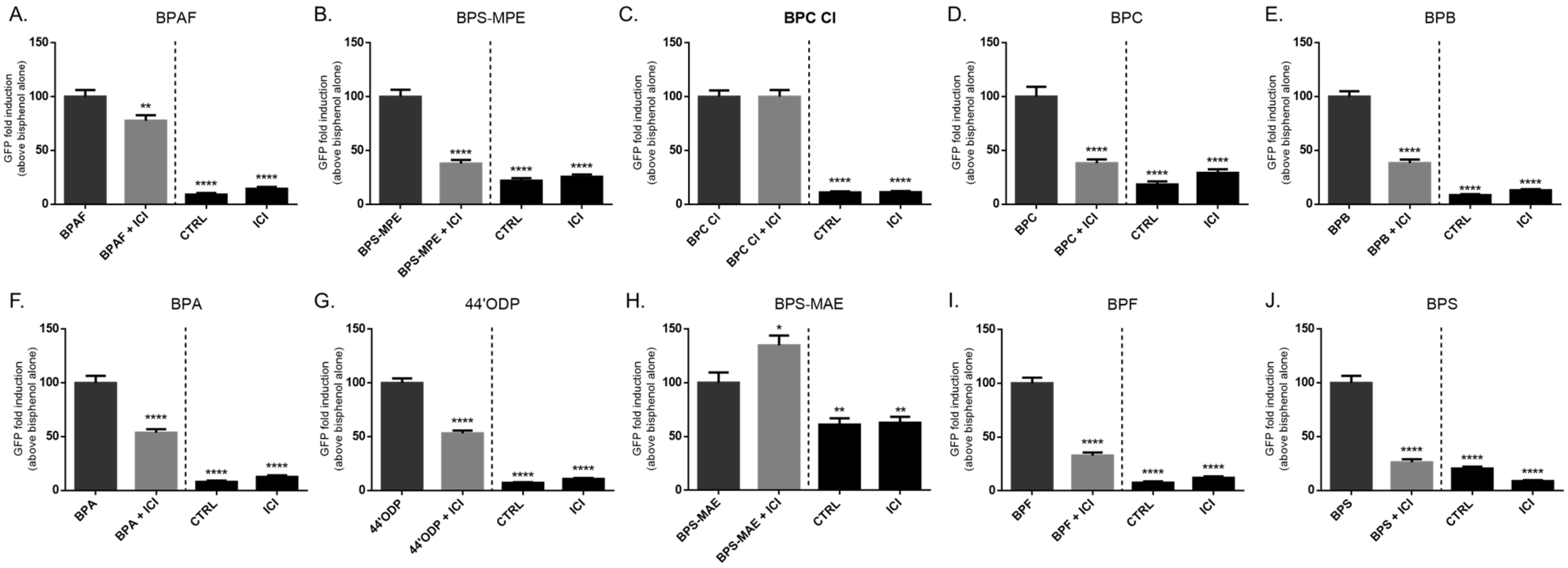


BPAF (0.34 mg/L; 1 µM), BPS-MPE (2 mg/L; 0.0059 µM), BPC (1 mg/L; 0.0039 µM), BPB (1.25 mg/L; 5.159 µM), BPA (5 mg/L; 22 µM), 44'ODP (1 mg/L; 0.005 µM), BPF (1 mg/L; 0.005 µM): 96h exposure with bisphenol and ICI 182,780 (1µM).

BPC Cl (0.3 mg/L; 0.0011 µM), BPS-MAE (0.625 mg/L; 0.0022 µM), BPS (100 mg/L; 0.4 µM): 48h exposure only with ICI 182,780 and from 48h exposure with bisphenol and ICI 182,780 until 96h. This doesn't work for BPC Cl and BPS-MAE even with the processing treatment.

The induction of GFP by bisphenols was determined as hundred percent of induction of GFP and then the decrease in GFP expression by co-exposure with ICI is expressed in fold of diminution.

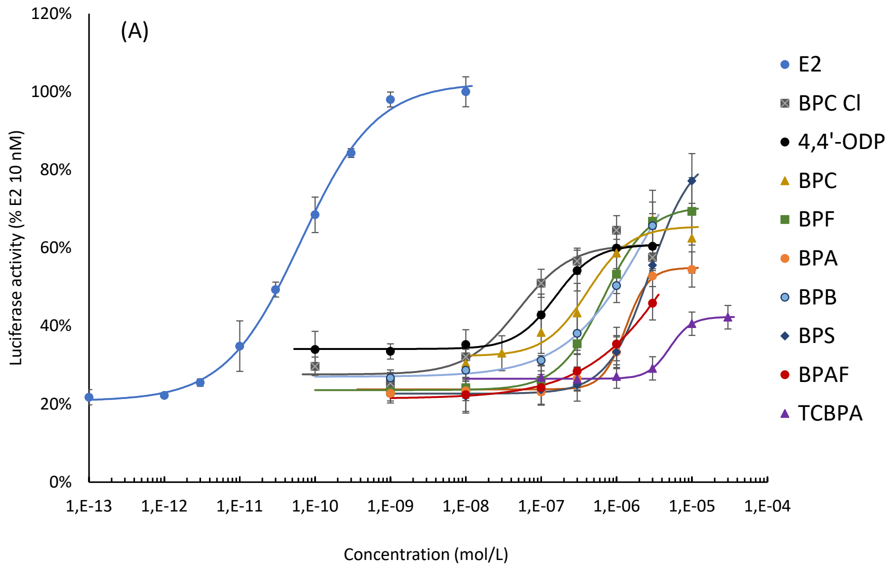


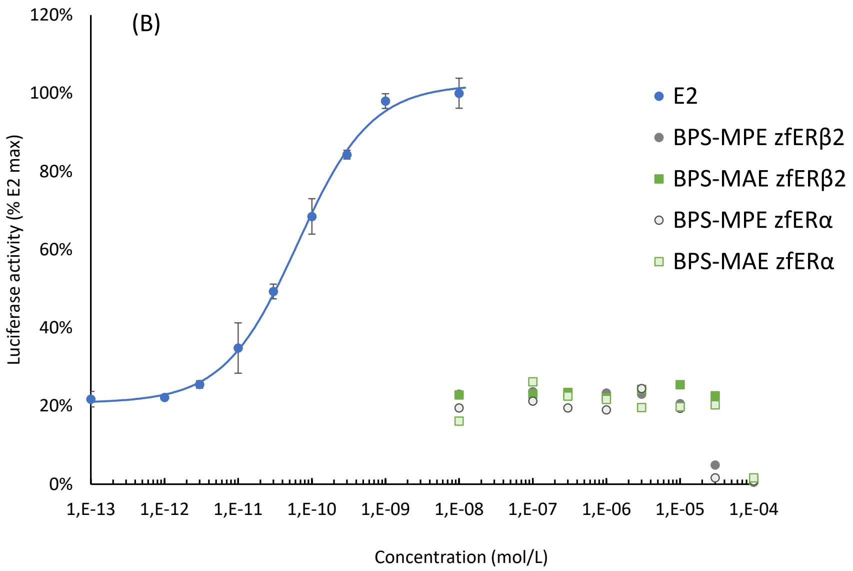


**Figure S7.** Induction of luciferase activity in the ZELH-zfERb2 cell line by bisphenols. (A) active BPs, (B) non active BPs. Data from 3 independent experiments were pooled to obtain the concentration-response curves.
